# Nutrient enrichment and herbivore exclusion disrupt the climate-driven balance between C_3_ and C_4_ plants

**DOI:** 10.64898/2026.08.17.744757

**Authors:** Joe Atkinson, Jodi N Price, Robert Buitenwerf, Nicholas G Smith, Ezinwanne Ezekannagha, Elizabeth T Borer, Cynthia Brown, Lars A Brudvig, Yvonne M Buckley, Miguel N Bugalho, Maria C Caldeira, Sofía Campana, Clinton Carbutt, Chris R Dickman, Ian Donohue, Nico Eisenhauer, Kenneth J Elgersma, Anu Eskelinen, Magda Garbowski, Sylvia Haider, Nicole Hagenah, Stanley Harpole, Yann Hautier, Anke Jentsch, Johannes MH Knops, Sally E Koerner, Mayank Kohli, Kimberly J Komatsu, Lauri Laanisto, Andrew DB Leakey, Petr Macek, Miaojun Ma, Andrew S MacDougall, Jason P Martina, Holly M Martinson, Rebecca L McCulley, John W Morgan, Meelis Pärtel, Steven C Pennings, Pablo L Peri, Sally A Power, Suzanne M Prober, Zhengwei Ren, Anita C Risch, Christiane Roscher, Mahesh Sankaran, Eric W Seabloom, Rachel Standish, Michelle Tedder, Risto Virtanen, Glenda M Wardle, Elizabeth F Waring, George Wheeler, Daniel M Griffith, Jens-Christian Svenning

## Abstract

The balance between plants with the C_3_ and C_4_ photosynthetic pathways shapes productivity, forage quality and carbon cycling in grasslands. Climate influences where each photosynthetic pathway is favoured, but how nutrient enrichment and changing herbivore communities alter this balance remains poorly understood. Across 3,186 plots at 112 sites on six continents, C_4_ dominance increases with the number of months in which mean maximum temperature reaches at least 23 °C with at least 20 mm of rainfall. In a globally replicated experiment, nutrient addition reduced C_4_ dominance, through both losses of C_4_ cover and gains in C_3_ cover, an effect driven by nitrogen. Herbivore exclusion alone had no effect, but combined with nutrient addition it produced the steepest declines in C_4_ dominance where C_4_ plants were initially dominant. Our findings show that nutrient enrichment can override the climatic advantage of C_4_ plants, with consequences for grassland productivity, grazing and carbon cycling.

## Introduction

Grasslands cover approximately 23% of the Earth’s land surface^1^ and vary widely in their dominance by C_3_ and C_4_ plants. Although only 3% of plant species globally exhibit the C_4_ pathway, C_4_-dominated vegetation accounts for approximately 20–25% of global terrestrial photosynthesis^2,3^. C_4_ grasses dominate tropical and subtropical savannas and many warm-temperate grasslands, where their photosynthetic efficiency under high temperatures and light, together with greater water and nutrient-use efficiencies, can support growth rates up to 40–50% higher than those of C_3_ species^4,5^. These characteristics of the C_4_ pathway help sustain productive grazing systems and contribute to fuel loads that support frequent fire regimes. C_3_ plants, in turn, dominate cooler, higher-latitude and high-elevation grasslands, remain active in the cool season when C_4_ plants are dormant, and can provide more digestible, protein-rich forage^6^. C_3_ forbs, including nitrogen-fixing legumes, also contribute significant and unique diversity in grasslands^7^.

With around 800 million people dependent on grasslands for their livelihoods^8^, predicting how the balance of C_3_–C_4_ plants in grasslands responds to global change is critical. Shifts in the C_3_– C_4_ balance could alter primary productivity, forage availability and quality, and carbon cycling with consequences for ecosystem functioning and food security. Recent work has documented the global decline of non-agricultural C_4_ grasses and increase in C_3_ species in response to increasing atmospheric CO ^2^. However, the relative influence of other global change factors including climate, nutrient availability, and herbivory, on C_3_ and C_4_ plant distributions remains poorly understood. Addressing these knowledge gaps is essential for predicting functional group responses to future environmental change.

The geographic distribution of C_3_ and C_4_ plants is governed primarily by climate^9–14^ and by disturbance regimes that maintain and promote open ecosystems^15,16^. Because C_4_ plants concentrate carbon dioxide (CO_2_) around Ribulose-1,5-bisphosphate carboxylase/oxygenase (RuBisCO), they minimise photorespiration and are generally more photosynthetically efficient than C_3_ plants under high temperature and low CO_2_^17^. The temperature crossover threshold is the point at which C_4_ photosynthesis becomes more advantageous than C_3_ photosynthesis due to greater efficiency under warm conditions. At recent historical atmospheric CO_2_ (∼300 ppm), C_4_ plants dominated in areas with mean annual temperatures above 22°C, and multiple monthly average maximum temperatures _≥_27°C, whereas C_3_ plants dominated below these thresholds^3,18^. Rising CO_2_ reduces photorespiration in C_3_ plants, effectively raising this crossover threshold and allowing C_3_ plants to compete in regions that were previously optimal for C_4_ species ^2^.

In addition to temperature (with C_4_ species commonly referred to as "warm-season grasses" and C_3_ as "cool-season grasses"), the timing and seasonality of precipitation are important determinants of C_3_–C_4_ dynamics^18,19^. Early work on Australian grass distributions demonstrated that C_4_ species are associated with hot, wet summers whereas C_3_ species favour cool, wet springs^19^. This pattern has since been supported by regional field studies, remote sensing analyses, and modelling approaches across the Northern Hemisphere ^2,13,18^. More recently, warm-season to cool-season rainfall ratios have been used to predict seasonal C_3_–C_4_ dynamics in Australia^20^. However, whether these relationships are consistent across grasslands globally remains unclear.

In addition to climate, local factors can strongly modify C_3_–C_4_ dynamics. Nitrogen availability may be a key regulator, as C_4_ plants are typically more efficient in nitrogen use and tend to outperform C_3_ plants under nitrogen-limited conditions^21–24^. Under elevated nitrogen, C_3_ plants may benefit relative to C_4_ plants, because C_3_ plants with RuBisCO-rich photosynthetic machinery can rapidly increase photosynthetic capacity, leaf area, and height, intensifying competition for light ^25^. This shift might be more pronounced in sites with initially high C_4_ abundance, where low-nutrient conditions structure plant communities. In such systems, nitrogen addition may effectively release C_3_ plants from strong limitation, leading to rapid competitive displacement of C_4_ species. This mechanism predicts that anthropogenic N inputs, now widespread in grasslands globally ^26^, may shift dominance from C_4_ to C_3_ plants. However, evidence of this shift is limited ^2,27^. Unlike nitrogen, there is no strong pathway-specific expectation for other macronutrients including phosphorus or potassium. However, productivity in many ecosystems is the result of co-limitation of multiple nutrients (most often nitrogen and phosphorus)^28,29^. Where phosphorus limits growth once nitrogen limitation is relieved, combined nutrient addition should increase biomass and canopy closure more than nitrogen alone. Therefore, phosphorus and nitrogen could affect C_3_–C_4_ balance through both independent and interactive effects on biomass and canopy dynamics.

Fire and herbivory also influence C_3_–C_4_ dynamics, although their effects may depend on climatic conditions and characteristics of the plant community^18,30^. C_4_ plants better tolerate biomass loss under favourable climatic conditions due to a higher photosynthetic quantum yield^31,32^. Fire has also been an important driver in the evolution and spread of C_4_-dominated grasslands^33,34^, with frequent burning potentially favouring C_4_ species by maintaining open canopies and high light availability^35^. Herbivory may produce similar effects by reducing vegetation biomass and opening the canopy. C_3_ plants generally have softer, more palatable tissue and slower regrowth following herbivory, whereas C_4_ plants are more resilient and recover faster following defoliation^6,36,37^. Consequently, changes in herbivore abundance could alter the competitive balance between C_3_ and C_4_ plants. For example, the loss of large herbivores may favour C_3_ plants in some grasslands, although such shifts can take decades to emerge^38^. Whether such shifts occur across grasslands globally remains largely untested.

Ultimately, climate determines where each photosynthetic pathway has a physiological advantage, while local conditions influence the extent to which this advantage is realised. Nutrient enrichment can alter competitive interactions by increasing biomass and canopy closure, while the removal of vertebrate herbivores can also release plant biomass from consumption and alter competitive interactions. Because these responses depend on initial community composition, they should be strongest where C_4_ plants are initially most dominant and strengthen over time.

We use data from a globally distributed, standardised, experimental network in grasslands (the Nutrient Network^39^) to test how climate, nutrients, and mammalian herbivory interact to shape the balance between C_3_ and C_4_ plants (Fig. 1). We first ask whether a temperature threshold, conditional on minimum precipitation requirements, explains C_4_ abundance across continents, and how it compares with other climatic variables and plot-level conditions. We hypothesise (H1) that C_4_ dominance is primarily determined by climate, with higher C_4_ proportions expected in regions with more months above the crossover threshold, and where annual mean temperature, warm-season rainfall, and ground-layer light levels are high (Fig. 1a). We then ask how local factors, including nutrients and herbivores, alter this balance, hypothesising (H2) that nutrient enrichment and herbivore exclusion influence C_4_ abundance (Fig. 1b). Specifically, we expect that (a) nutrient addition, particularly nitrogen, will reduce the competitive advantage of C_4_ plants and favour C_3_ plants; (b) herbivore exclusion will favour C_3_ species, as herbivores preferentially feed on C_3_ plants and promote open, high-light conditions; (c) the strongest declines will occur when nutrient addition and herbivore exclusion are applied together; and (d) the effects of nutrient addition and herbivore exclusion will correspond with reduced ground-layer light, increased treatment duration, and will depend on initial C_4_ cover.

**Figure 1.**
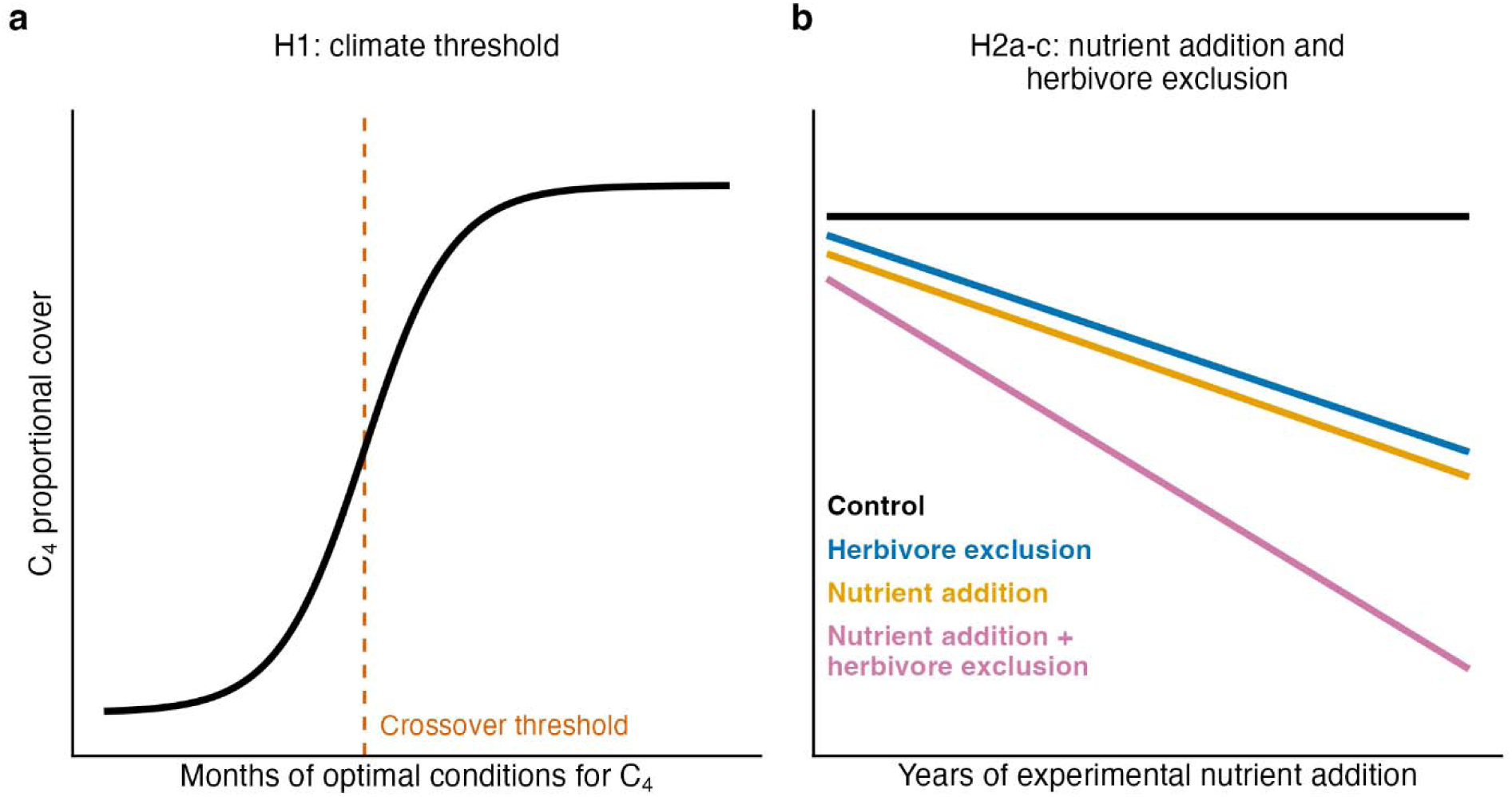
Conceptual framework and hypotheses. a, H1: C_4_ dominance increases sharply once the number of months that are both warm and wet exceeds a crossover threshold. b, H2a–d: the predicted effects of nutrient addition and herbivore exclusion (fencing) and both combined. Nutrient addition, particularly nitrogen, and herbivore exclusion are expected to reduce the competitive advantage of C_4_ plants and promote C_3_ plants, with the strongest effects expected when combined and to strengthen over time. Not shown schematically initial dominance and light components of H2d: declines should be strongest where C_4_ plants were initially most dominant and where ground-layer light is most strongly reduced.

In this work, we show that climate is a strong predictor of C_4_ dominance across grasslands globally, while experimental manipulations reveal how nutrient availability and herbivory can alter the balance between C_3_ and C_4_ plants locally. Nitrogen addition drives declines in C_4_ dominance, through both losses of C_4_ cover and gains in C_3_ cover. Excluding herbivores has no effect on C_3_–C_4_ balance on its own, but when combined with nutrient addition it produces the greatest declines in C_4_ dominance, particularly at sites where C_4_ plants were initially dominant.

## Results

### Climate is a key determinant of C_3_–C_4_ abundance

Our findings support H1. The number of months with a mean maximum temperature ≥ 23**°**C and ≥20 mm of precipitation was strongly associated with C_4_ proportional cover (Table 1). Averaged across sites, predicted C_4_ cover rose from about 1% at sites with no qualifying months to 19% with four, 66% with seven and over 94% with nine or more (Fig. 2). This was the best supported individual threshold of the 110 thresholds tested (Supplementary Table 1) and a better predictor than mean annual temperature, mean maximum temperature, and the warm- to cool-season rainfall ratio (Supplementary Table 6). Temperature and rainfall variables together explained slightly more variation in C_4_ cover than the threshold alone, but the two approaches captured largely overlapping information. When modelled alongside other climate variables, the threshold retained a significant, although modest, independent effect on C_4_ cover (p = 0.041; Supplementary Table 6), while maximum temperature of the warmest month and the ratio of warm- to cool-season rainfall also remained significant (Supplementary Table 6; Supplementary Fig. 5). This indicates that the threshold captures a climate signal that overlaps with, but is not fully explained by, these other variables. Its effect was robust to sampling year, the inclusion of mean annual precipitation, and accounting for spatial covariance among sites (Supplementary Tables 3 and 4). Furthermore, at the 86 sites where both were available, the threshold explained more variation in C_4_ cover than the satellite-derived C_4_ distribution of Luo et al.^2^ (marginal R^2^ 0.52 vs 0.27; Supplementary Table 2).

**Figure 2.**
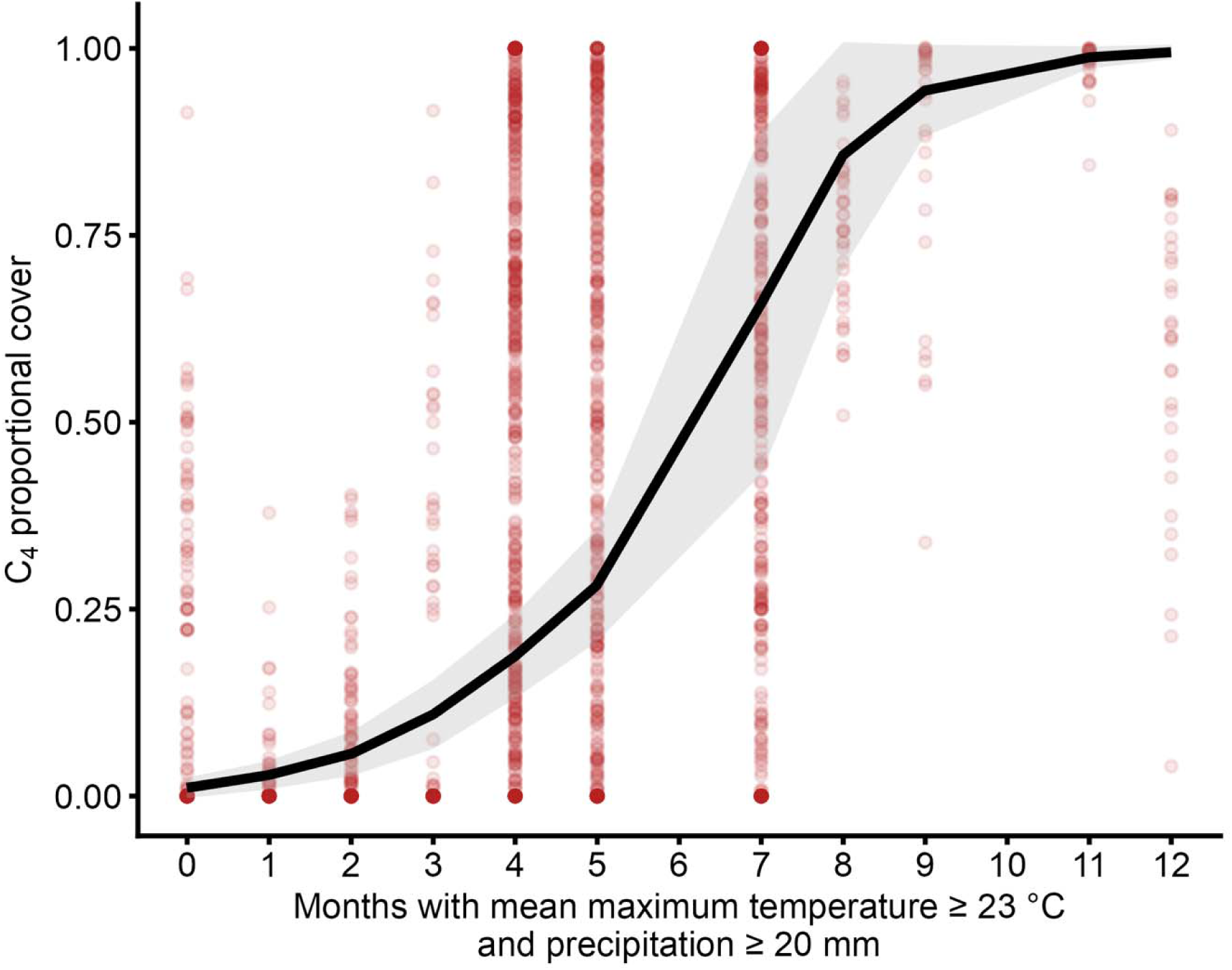
C_4_ dominance rises steeply with the number of warm, wet months. Proportional C_4_ cover in relation to the number of months in which mean maximum temperature was ≥23 °C and precipitation was ≥20 mm (the best-supported crossover threshold; Supplementary Table 1). Points represent observed values for individual plots (3,186 plots across 112 sites). The line shows predictions from the ordered beta model in Table 1, averaged across sites, and shading indicates 95% confidence intervals.

**Table 1.** Climatic threshold and ground-level light as predictors ofC_4_ dominance.

| Predictor | Odds ratio | 95% CI | z | p |
|---|---|---|---|---|
| Intercept | 0.002 | 0.001–0.006 | −10.44 | <0.001 |
| Months ≥23 °C and ≥20 mm (per SD) | 113.19 | 41.12–311.53 | 9.15 | <0.001 |
| Proportion of PAR reaching ground level (per SD) | 1.00 | 0.89–1.12 | −0.02 | 0.982 |
| Random-effect variance: site | 20.64 |  |  |  |
| Random-effect variance: block within site | 0.27 |  |  |  |
| Observations (plots) | 3,186 |  |  |  |
| Sites | 112 |  |  |  |
| Marginal R<sup>2</sup> | 0.42 |  |  |  |
| Conditional R<sup>2</sup> | 0.82 |  |  |  |

Ordered beta mixed model of proportional C<sub>4</sub> cover (C<sub>4</sub> cover divided by combined C<sub>3</sub> and C<sub>4</sub> cover) in 3,186 plots at 112 sites, with random intercepts for site and for block nested within site. Continuous predictors were standardised (mean = 0, SD = 1), so odds ratios represent changes per SD increase in the predictor. The intercept represents the model estimate when both predictors are at their mean values. CI, confidence interval; PAR, photosynthetically active radiation; SD, standard deviation; z, Wald statistic.

Previously reported thresholds of mean monthly maximum temperature thresholds of ≥27**°**C and ≥25 mm of precipitation produced similar patterns but were less strongly supported (ΔAIC = 19.9; Supplementary Table 1). Despite the clear overall pattern, C_4_ dominance varied substantially among sites with the same number of qualifying months, particularly at four, five, and seven months (Fig. 2). This variation indicates that climate alone does not explain differences in C_4_ dominance among sites. Local-scale factors—such as nutrient availability, herbivory, or other site-level conditions—likely modulate the expression of C_4_ dominance under favourable climatic conditions.

In addition to the crossover threshold, the log ratio of warm-season to cool-season precipitation was a significant predictor of C_4_ dominance across all plots (Supplementary Fig. 5), with higher values (greater warm-season rainfall relative to cool-season rainfall) associated with greater C_4_ dominance. Hence, C_4_ plants benefit from water availability coinciding with their peak growing season. Beyond climate, we included the proportion of photosynthetically active radiation (PAR) reaching the ground at the plot level as a measure of local canopy openness, and this was not related to C_4_ cover (Table 1). Higher C_4_ dominance at sites with more months above the climate threshold was associated primarily with lower proportional cover of C_3_ grasses and, to a lesser extent, C_3_ forbs (Supplementary Table 5).

### Nutrient addition reduces C_4_ dominance

While the climate model (H1) identified strong support for the crossover threshold, it explained less than half of the variation in C_4_ dominance (marginal R² = 0.42; Table 1), indicating that other factors also influenced C4 abundance. We next tested how nutrient enrichment and herbivore exclusion influenced C_4_ abundance within the climatic conditions that support C_3_–C_4_ coexistence. At the mean experiment duration (5.4 years) and mean initial C_4_ cover, C_4_ cover was lower in NPK (odds ratio 0.61, 95% CI 0.50-0.75) and NPK+Fence (0.53, 0.42-0.67) plots than in control plots, whereas fencing alone had no detectable effect (0.92, 0.73-1.16; Table 2, Fig. 3). Fencing alone modestly reduced the proportion of PAR reaching the ground (from 0.34 to 0.28; odds ratio 0.76, 95% CI 0.65-0.88), while NPK and NPK+Fence reduced it further, to 0.23 and 0.18 respectively (Supplementary Table 14).

**Figure 3.**
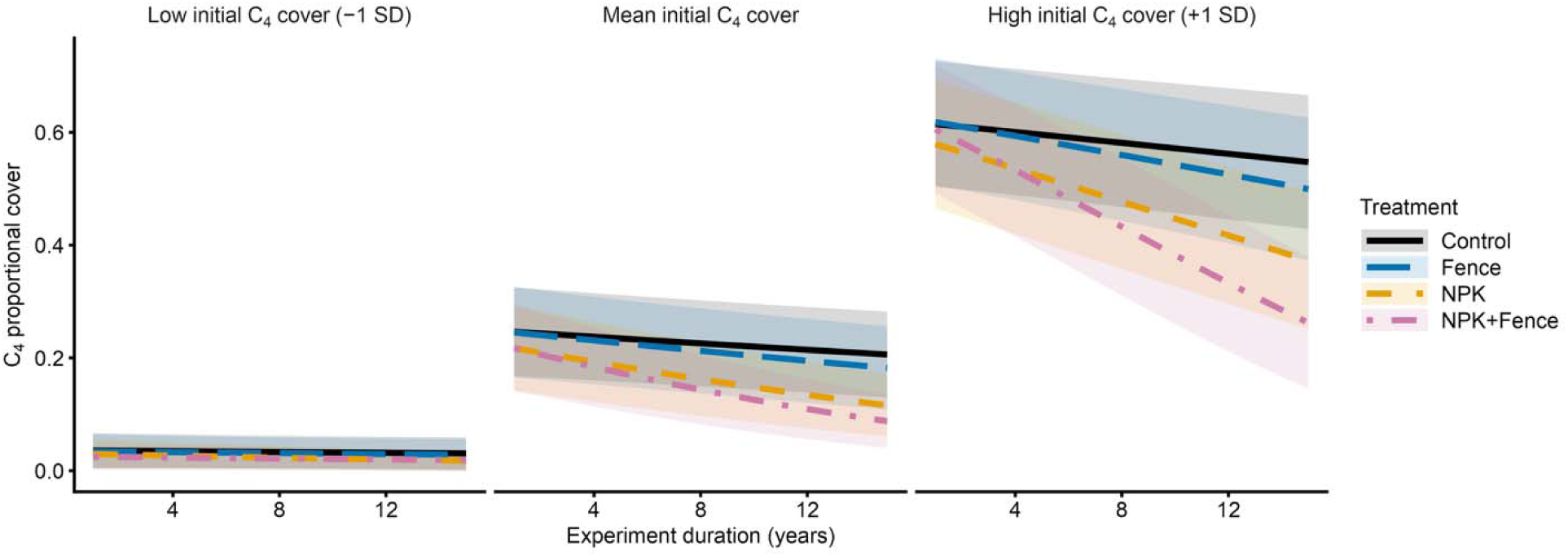
Declines in C_4_ dominance were greatest under combined nutrient addition and vertebrate herbivore exclusion, particularly where C_4_ plants were initially dominant. Predicted proportional C_4_ cover over the experiment duration for each treatment at low (-1 SD), mean and high (+1 SD) initial C_4_ cover, based on the ordered beta model including initial C_4_ cover (Table 2, model 2). Predictions are averaged across sites and shading shows 95% confidence intervals. Conditional slopes are provided in Supplementary Table 8, and the number of sites contributing at each plot age is given in Supplementary Table 7. Control = unfenced plots without added nutrients; Fence = herbivore exclusion; NPK = nitrogen, phosphorus and potassium addition.

**Table 2.** Effects of nutrient addition and herbivore exclusion on C_4_ dominance over time.

| Predictor | Odds ratio, model 1 | 95% CI, model 1 | p, model 1 | Odds ratio, model 2 | 95% CI, model 2 | p, model 2 |
|---|---|---|---|---|---|---|
| Intercept | 0.12 | 0.04–0.40 | <0.001 | 0.14 | 0.07–0.28 | <0.001 |
| Fence | 0.91 | 0.73–1.14 | 0.423 | 0.92 | 0.73–1.16 | 0.485 |
| NPK | 0.58 | 0.48–0.70 | <0.001 | 0.61 | 0.50–0.75 | <0.001 |
| NPK+Fence | 0.51 | 0.40–0.64 | <0.001 | 0.53 | 0.42–0.67 | <0.001 |
| Plot age | 0.97 | 0.95–0.98 | <0.001 | 0.97 | 0.95–0.99 | 0.010 |
| Fence × plot age | 0.98 | 0.95–1.01 | 0.212 | 0.98 | 0.95–1.02 | 0.317 |
| NPK × plot age | 0.94 | 0.91–0.96 | <0.001 | 0.95 | 0.92–0.98 | <0.001 |
| NPK+Fence × plot age | 0.90 | 0.87–0.93 | <0.001 | 0.92 | 0.89–0.95 | <0.001 |
| Initial C<sub>4</sub> cover (per SD) | – | – | – | 18.08 | 9.05–36.11 | <0.001 |
| Fence × initial C<sub>4</sub> cover | – | – | – | 0.98 | 0.77–1.25 | 0.883 |
| NPK × initial C<sub>4</sub> cover | – | – | – | 0.87 | 0.70–1.08 | 0.218 |
| NPK+Fence × initial C<sub>4</sub> cover | – | – | – | 0.87 | 0.68–1.11 | 0.249 |
| Plot age × initial C<sub>4</sub> cover | – | – | – | 0.99 | 0.97–1.01 | 0.261 |
| Fence × plot age × initial C<sub>4</sub> cover | – | – | – | 0.99 | 0.95–1.02 | 0.464 |
| NPK × plot age × initial C<sub>4</sub> cover | – | – | – | 0.98 | 0.95–1.01 | 0.163 |
| NPK+Fence × plot age × initial C<sub>4</sub> cover | – | – | – | 0.93 | 0.89–0.96 | <0.001 |
| Random-effect variance: site | 15.92 |  |  | 5.15 |  |  |
| Random-effect variance: block within site | 0.21 |  |  | 0.22 |  |  |
| Random-effect variance: plot within block | 0.43 |  |  | 0.43 |  |  |
| Observations (plot-years) | 4,179 |  |  | 4,179 |  |  |
| Sites | 48 |  |  | 48 |  |  |
| Marginal R<sup>2</sup> | 0.01 |  |  | 0.55 |  |  |
| Conditional R<sup>2</sup> | 0.95 |  |  | 0.94 |  |  |

Ordered beta mixed models of the proportion of C<sub>4</sub> cover in 4,179 plot-years at 48 sites, without (model 1) and with (model 2) initial C<sub>4</sub> cover, with random intercepts for site, block within site and plot within block. Treatments are compared with unfenced control plots without added nutrients. Plot age (years since treatments began) was mean-centred, and initial C<sub>4</sub> cover (site mean proportion in the pre-treatment survey) was mean-centred and scaled by its standard deviation. Thus main effects are evaluated at the mean plot age and, in model 2, at the mean initial C<sub>4</sub> cover. Odds ratios for initial C<sub>4</sub> cover terms are per SD (0.33). Dashes indicate terms not included in model 1. NPK, nitrogen, phosphorus and potassium addition; Fence, herbivore exclusion; CI, confidence interval, SD, standard deviation.

Analysis of individual nutrient effects identified nitrogen as the main driver of C_4_ decline (Table 3, Fig. 4). Nitrogen addition reduced C_4_ cover in the factorial model (estimate = -0.07, 95% CI - 0.12 to -0.01, p = 0.020; Supplementary Table 15). The largest declines occurred when N was combined with P (NP -0.08, NPK -0.10, and NPK+Fence -0.09; Table 3), although these responses did not differ significantly from N alone in pairwise comparisons (Supplementary Table 16). P and K alone had no detectable effects. The factorial model also showed no significant N × P interaction (estimate = -0.06, p = 0.12), indicating no evidence that the combined effect of N and P was greater than expected from their individual effects.

**Figure 4.**
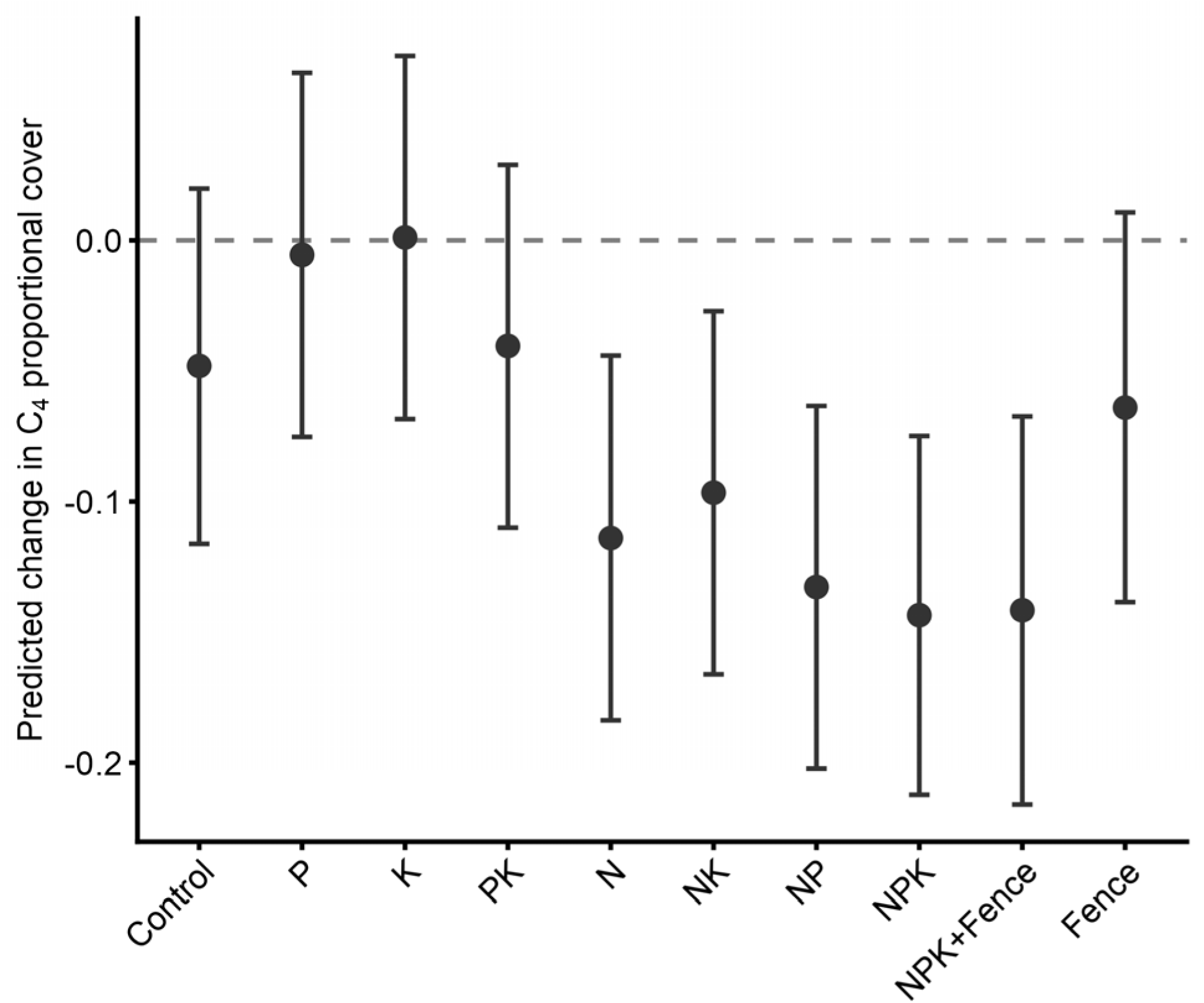
Nitrogen-containing treatments reduced C4 dominance after four years of application. Predicted change in proportional C4 cover between the pre-treatment survey and year 4 for each treatment, based on the linear mixed model in Table 3 (836 plots at 27 sites). Points show model predictions and error bars show 95% confidence intervals. The dashed line indicates no change and negative values indicate declines in C4 cover. Pairwise comparisons between treatments are provided in Supplementary Table 16. N = nitrogen; P = phosphorus; K = potassium; Fence = herbivore exclusion.

**Table 3.** Effects of individual nutrients and fencing on the change in C_4_ dominance after four years of treatment application.

| Treatment | Estimate | 95% CI | t | p |
|---|---|---|---|---|
| Intercept (control) | −0.05 | −0.12 to 0.02 | −1.39 | 0.165 |
| P | 0.04 | −0.01 to 0.10 | 1.48 | 0.140 |
| K | 0.05 | −0.01 to 0.11 | 1.71 | 0.087 |
| PK | 0.01 | −0.05 to 0.06 | 0.27 | 0.790 |
| N | −0.07 | −0.12 to −0.01 | −2.27 | 0.023 |
| NK | −0.05 | −0.10 to 0.01 | −1.68 | 0.092 |
| NP | −0.08 | −0.14 to −0.03 | −2.94 | 0.003 |
| NPK | −0.10 | −0.15 to −0.04 | −3.38 | 0.001 |
| NPK+Fence | −0.09 | −0.16 to −0.03 | −2.93 | 0.004 |
| Fence | −0.02 | −0.08 to 0.05 | −0.49 | 0.623 |
| Random-effect variance: site | 0.02 |  |  |  |
| Random-effect variance: block within site | 0.004 |  |  |  |
| Observations (plots) | 836 |  |  |  |
| Sites | 27 |  |  |  |
| Marginal R<sup>2</sup> | 0.04 |  |  |  |
| Conditional R<sup>2</sup> | 0.42 |  |  |  |

The decline in proportional C_4_ cover under nutrient addition reflected both a reduction in absolute C_4_ cover and an increase in absolute C_3_ cover. Relative to control plots, absolute C_4_ cover was lower under NPK (cover ratio 0.74, 95% CI 0.60-0.91) and NPK+Fence (0.70, 0.55-0.89) and declined more rapidly over time (-5.8% and -7.7% per year, compared with -2.0% in controls). Under these same treatments, absolute C_3_ cover was higher (NPK 1.24, 1.16-1.33; NPK+Fence 1.32, 1.22-1.44) and increased over time (+1.1% and +1.9% per year respectively). Fencing alone affected neither C_3_ nor C_4_ cover (Supplementary Tables 12 and 13). Among C_3_ groups, C_3_ grasses increased under NPK (treatment × year 0.030, p = 0.014; adjusted p = 0.043), whereas C_3_ forbs increased only under NPK+Fence (0.048, p < 0.001; adjusted p < 0.001). Neither group responded to fencing alone (Supplementary Table 18; Supplementary Fig. 7). C_3_ woody cover showed no treatment response after correction for multiple comparisons, consistent with its low cover at most sites.

### Initial community state and time

Initial C_4_ cover was the strongest single predictor of proportional C_4_ cover, and including it in the model substantially improved explanatory power (marginal R^2^ 0.55 vs 0.01; Table 2). Sites with higher initial C_4_ proportional cover tended to retain relatively higher proportions over time. C_4_ cover also declined over time in control plots, although this was detectable only at mean and high initial C_4_ cover. At low initial C_4_ cover (-1 SD), control plots showed no detectable temporal change (odds ratio per year 0.99, 95% CI 0.95-1.02, p = 0.48). At mean initial cover, C_4_ cover declined slightly (0.97, 0.95-0.99, p = 0.010), while at high initial cover (+1 SD) the decline was also detectable (0.96, 0.94-0.98, p < 0.001; Supplementary Table 8).

### Initial C_4_ dominance and time modify treatment effects

Nutrient addition accelerated declines in C_4_ dominance over time. At mean initial C_4_ cover, C_4_ proportional cover declined faster than in control plots under NPK (odds ratio 0.95, 95% CI 0.92-0.98) and NPK+Fence (0.92, 0.89-0.95), but not under fencing alone (0.98, 0.95-1.02; Table 2).

However, the extent to which initial C_4_ cover modified treatment effects on the rate of decline differed between nutrient treatments. The significant three-way interaction between NPK+Fence, plot age and initial C_4_ cover (odds ratio 0.93, 0.89-0.96, p < 0.001) indicates that the additional decline relative to controls became more pronounced as initial C_4_ cover increased. In contrast, under NPK alone, the additional decline relative to controls did not vary significantly with initial C_4_ cover (NPK × plot age × initial C_4_ cover, 0.98, 0.95-1.01, p = 0.16). At low initial C_4_ cover, C_4_ dominance declined only under NPK (0.96 per year, p = 0.042). At high initial C_4_ cover, it declined under all treatments, with the largest decline under NPK+Fence (Supplementary Table 8; Fig. 3).

### Exploratory analyses of C_4_ grass lineages

In exploratory analyses, we partitioned the overall C_4_ response by functional group and grass lineage (Supplementary Fig. 8, Supplementary Tables 20 and 21). Estimates for C_4_ grasses alone were similar to those for all C_4_ vegetation (Supplementary Table 21), while C_4_ forbs, which occurred at only eight sites, contributed little to the overall response. Within grasses, declines were concentrated in the lineage Andropogoneae (18 sites), which declined under all five nitrogen-containing nutrient addition treatments (adjusted p ≤ 0.008) but showed little response to P, K or fencing alone. Chloridoideae (22 sites), the MPC clade (13 sites) and *Aristida* (8 sites) showed no significant response to any treatment after correction for multiple comparisons. The MPC clade tended to increase under NPK+Fence (raw p = 0.009; adjusted p = 0.054). A model comparing lineages at the 22 sites where at least two co-occurred with a treatment × lineage interaction confirmed that treatment responses differed among lineages (Supplementary Table 22). However, because the lineage-specific models were fitted to different subsets of sites, these results should be interpreted cautiously.

## Discussion

Our results demonstrate that climate is a key driver of C_4_ dominance, particularly the coincidence of warm temperatures and sufficient rainfall. The strongest predictor was the number of months in which both temperature and precipitation exceeded the crossover criteria, highlighting the importance of the duration of favourable conditions. The seasonal distribution of rainfall was also a strong predictor, indicating that the timing of precipitation relative to growing-season temperatures provides additional explanatory power. Sites may therefore experience sufficient warm months to exceed the thermal crossover threshold but still not support C_4_ dominance if rainfall is concentrated in cooler months^40^. The findings underscore the importance of considering both temperature and the seasonal distribution of rainfall when predicting vegetation responses to climate change.

Our best-supported crossover threshold (monthly maximum temperature average of 23 °C) is lower than theoretical predictions (∼30 °C at current CO_2_ levels^11^) and some regional estimates (27 °C^18^) but closely aligns with the global estimates using monthly mean temperatures^3^ (22 °C). Importantly, several models with crossover temperatures near 23 °C were also well supported, suggesting that a single universal threshold is unlikely. We also observed substantial variation in C_4_ proportional cover around the estimated threshold, indicating that other factors beyond temperature and precipitation contribute to C_4_ distributions. The crossover concept is based on differences in leaf-level quantum yield between C_3_ and C_4_ plants. Our simple count-based metric attempts to capture the advantage of the C_4_ pathway over the year: each month in which temperatures exceed the crossover and rainfall supports growth potentially extends the period over which C_4_ plants can “out-assimilate” C_3_ neighbours, and this greater carbon gain should then translate into faster growth and biomass accumulation^4^. Consistent with this interpretation, thresholds requiring warm and wet conditions in the same month were much better supported than models counting warm and wet months separately (ΔAIC ≥ 21; Supplementary Table 1). The steep rise in C_4_ dominance between about four and eight qualifying months (Fig. 2) is consistent with a cumulative advantage of C_4_ plants as the duration of favourable conditions increases, potentially overcoming the earlier seasonal growth of C_3_ plants.

Classical crossover models were originally developed for grasses, where growth forms are more comparable and differences associated with photosynthetic pathway can be more directly isolated^10,12^. In contrast, our observational dataset also includes non-grass C_3_ and C_4_ species, introducing additional variation that is not captured in classic pathway-based models. Variation in C_4_ cover around the estimated threshold may therefore reflect differences in growth form, species composition and other site-level environmental or ecological factors. However, we cannot distinguish these influences from limitations of the threshold metric itself, particularly given the heterogeneity in species composition across our global dataset.

Climate change is likely to affect the C_3_–C_4_ balance through multiple interacting pathways. First, rising temperatures and changes in precipitation will alter where (and for how much of the year) conditions favour C_4_ photosynthesis. As our results demonstrate, additional warm months will only favour C4 plants where sufficient rainfall also supports growth; warming accompanied by increased drought may therefore provide little advantage. Second, rising CO_2_, will interact with temperature and water availability to influence the relative performance of C_3_ and C_4_ plants^10,12^. Elevated CO_2_ suppresses photorespiration in C_3_ plants, increasing their quantum yield and raising the predicted crossover temperature^10^, with potentially greater benefits in warmer climates. However, C4 plants also respond to elevated CO_₂_ through reduced stomatal conductance and water loss, potentially gaining an advantage where water is limiting^41^. Therefore, rising CO_2_ will not necessarily reduce C_4_ dominance; its effect will depend on the relative benefits to each pathway under prevailing temperature and water conditions, and how they change together through time^11^.

Ground-level light did not predict C_4_ dominance across sites. Because ground-level PAR was measured beneath the canopy, it reflects vegetation structure and may itself be influenced by plant composition, making it difficult to infer a causal relationship with C_4_ abundance from the observational data. However, the experimental results provide insight into how vegetation structure and light availability respond to nutrient addition and herbivore exclusion. Nutrient addition reduced the proportion of light reaching the ground by about a third relative to unfertilised, unfenced controls, and by nearly half when combined with fencing (Supplementary Table 14). These changes coincided with reductions in C_4_ dominance, including at climatically favourable sites. Fencing alone reduced ground-level light only modestly and did not reduce C_4_ dominance. Other disturbances, such as fire, which we did not examine, may also maintain open, high-light conditions and thereby favour C_4_ species in many systems^35^.

Overall, both warming and rising CO_2_ – central aspects of human-induced climate change – are likely to strongly influence the structure of future grassland ecosystems^42^. However, our climate models explained less than half of the variation in C_4_ dominance, while our experiments show that local drivers can substantially alter the C_3_–C_4_ balance within a given climate.

Although climate defines the broad distributional template, our experimental results demonstrate that local drivers can substantially restructure C_3_–C_4_ dynamics within that climatic context. Nutrient addition consistently reduced the proportion of C_4_ plants, both independently and in combination with herbivore exclusion, whereas herbivore exclusion alone had little effect. Hence, nutrient enrichment, especially nitrogen addition, can reduce C_4_ dominance, even in climates otherwise favourable for their persistence.

Nutrient addition is well known to reduce plant diversity by intensifying competition for light, with nitrogen consistently identified as the primary driver of grassland species loss across regional and global scales^43–45^. Early experimental work in Minnesota grasslands demonstrated that nitrogen addition favoured early-successional C_3_ forbs and cool season grasses over late-successional warm season C_4_ grasses through competitive exclusion under closed canopies^45,46^. Our results are broadly consistent with this mechanism showing that nutrient enrichment drives a systematic decline in C_4_ cover and a corresponding increase in C_3_-dominated components of the community, although the identity of the plant groups responding varied with treatment.

Nutrient-driven declines in C_4_ dominance accumulated over time, and the temporal effect of nutrient addition combined with herbivore exclusion was strongest where C_4_ plants were initially dominant (Table 2). At initially C_4_-dominated sites, nutrient addition combined with herbivore exclusion reduced C_4_ cover from 55% to 23% over 14 years. These substantial reductions occurred in communities initially highly favourable to C_4_ plants demonstrating how strongly nutrient enrichment can destabilise ecosystems otherwise suited to C_4_ plants. Because the strongest declines occurred at sites where C_4_ grasses were initially dominant, our results suggest that C_4_-dominated ecosystems may be disproportionately vulnerable to anthropogenic nutrient loading. The nutrient addition rate in our experiment (10 g m^−2^ y^−1^) was considerably higher than most atmospheric deposition rates worldwide^47^, although lower atmospheric inputs have also been implicated in other continental-scale, ecosystem-level changes ^48,49^. Our results therefore point to potentially greater risks where nutrient inputs are high, such as fertilised or runoff-affected grasslands and areas experiencing severe atmospheric deposition^50^. However, because we applied a single, relatively high rate of nitrogen addition, we cannot determine whether low chronic inputs such as atmospheric deposition produce proportional effects. Dose-response experiments and long-term monitoring over deposition gradients are needed to address this question. Determining whether such shifts are already underway represents an important next step for global-change research.

The identity of the limiting nutrient also varies among grasslands, with P-limitation more common in the tropics and subtropics^51^. Across our sites, declines in C_4_ proportional cover under NPK+Fence were greater where soil P was higher (Supplementary Table 17). This suggests that soil P availability may modify the response to combined nutrient addition and herbivore exclusion, although we found no evidence that responses to P addition or NPK alone varied with soil P. However, our fully factorial experimental data (27 sites) included only one tropical site, preventing a rigorous comparison of responses to different nutrient combinations between tropical savannas and other biomes.

We found no overall main effect of herbivore exclusion on C_4_ dominance, although its effects depended on NPK addition, initial C_4_ abundance, and plot age. Herbivores are often expected to favour C_4_ plants by maintaining open, high-light conditions. However, fencing alone reduced ground-level light by only 18%, compared with 32% under NPK, and did not increase C_3_ cover. This suggests that herbivore exclusion alone did not consistently alter vegetation structure sufficiently to shift the balance between C_3_ and C_4_ plants over the study period. In contrast, combining nutrient addition with herbivore exclusion produced the lowest ground-level light (47% below unfertilised, unfenced controls) and the largest declines in C_4_ dominance, consistent with an interaction between nutrient enrichment and reduced herbivory in promoting vegetation closure. Previous work in the Nutrient Network has shown that herbivore effects on plant communities depend partly on whether herbivores alter ground-level light^52^, as well as on soil nutrients and rainfall^53^. However, herbivore regimes in unfenced plots were not standardised and instead reflected ambient herbivore composition and densities. Given that contemporary herbivore populations are often substantially lower than historical levels worldwide^54^, grazing pressure may have been insufficient at some sites to substantially alter canopy structure.

Herbivory effects on vegetation can be complex and context dependent. Some studies report that grazing promotes C_4_ plants under warming^55^, while others show reductions or variable responses in C_4_ biomass with grazing ^56^. Consistent with our results, climate can override the effect of grazing on C_4_ abundance^57^. Importantly, herbivores do not select plants according to photosynthetic pathway per se, but according to traits such as palatability, growth form, and phenology, which may be associated with different photosynthetic pathways^58^. Exclosure experiments also assess the response to the removal of herbivory, rather than the effects of herbivory itself. Consequently, they may not capture longer-term effects of grazing or the legacies of previous herbivore activity^59^. Although we found little overall effect of herbivore exclusion, herbivory remains a plausible driver of C_3_–C_4_ dynamics whose effects depend on climate, vegetation characteristics and grazing history^60,61^.

In control plots, C_4_ cover declined with experimental duration in sites with moderate to high initial C_4_ cover. This is in concordance with the results of Luo et al.^2^, who reported a global decrease in the extent of natural C_4_ grasslands from 15% to 14.2% between 2001 and 2019 (approximately 1 million km^2^), which they attributed to rising atmospheric CO_2_. Increasing CO_2_ reduces the relative photosynthetic advantage of C_4_ plants, from an estimated 70% greater efficiency at 200 ppm (Last Glacial Maximum) to 20% at 500 ppm, and almost zero at 700 ppm^10^. Consistent with a broader temporal influence, C_4_ cover in control plots also declined with calendar year across climatic contexts (Supplementary Table 9). Rising CO_2_ may have contributed to this decline, although its effects on the relative performance of C_3_ and C_4_ plants depend on temperature and water availability^41,42^. Other factors that have changed over this period, including atmospheric nitrogen deposition, may have also contributed.

In addition to CO_2_ effects, nutrient-driven increases in C_3_ biomass in fertilized plots may have elevated propagule pressure into adjacent control plots^62^, indirectly contributing to declines in C_4_ cover. Atmospheric nitrogen deposition may be a contributing factor to these declines, given our experiment showed a strong reduction of C_4_ plants under added nitrogen. However, these patterns are not universal with other literature providing contrasting evidence. For example, increases in C_4_ dominance have been documented across eastern Australia over the past 15 years, where seasonal rainfall has shifted to a more summer-dominant rainfall regime ^20^. We encourage further examination of long-term vegetation monitoring data to test the generality of a decline in C_4_ plants aligning with these putative drivers.

When considering grasses specifically, declines in C_4_ cover under nutrient addition were concentrated in Andropogoneae, which declined significantly under all nitrogen-containing treatments, whereas Chloridoideae showed no detectable response. No other lineage responded significantly after correction for multiple comparisons, although the MPC clade, dominated at our sites by *Panicum* and *Setaria*, tended to increase under NPK combined with fencing. Andropogoneae are the dominant C_4_ lineage across mesic tropical and subtropical grasslands and savannas globally, and their cover and functional dominance have been linked to fire regimes, rainfall seasonality, and low soil nitrogen availability^18,63,64^. The nitrogen-specificity of the response (with no equivalent effect of phosphorus or potassium) is consistent with evidence that Andropogoneae tend toward conservative nutrient-use strategies^65^ and corresponds with the dominance of this lineage in higher-rainfall, dystrophic savannas. At least one study suggested that some perennial Andropogoneae may stimulate nitrogen leaching rather than retaining added nitrogen^66^. However, our general prediction for N-driven losses in C_4_ species was based on the premise that the increased nitrogen-use efficiency of the pathway should mean they lose competitive advantage regardless of lineage. The concentration of declines in Andropogoneae therefore suggests that responses to nutrient enrichment may depend on taxonomic identity as well as photosynthetic pathway. We propose this as a hypothesis for testing in future experiments designed around C_4_ grass lineages.

The identity of the functional groups replacing declining C_4_ grasses also depended on herbivore exclusion. C_3_ grasses increased under NPK alone, while C_3_ forbs increased under NPK combined with herbivore exclusion. This difference may reflect an influence of herbivores on the functional groups able to otherwise capitalise on the competitive space opened by declining Andropogoneae. For example, herbivores may have suppressed broadleaved competitors in unfenced plots, allowing C3 grasses to increase under nutrient addition alone. However, grazer identity and density varied among sites, and the mechanisms underlying these contrasting responses remain unclear^67,68^. These patterns across functional groups and lineages are based on subsets of sites, as not all sites contained all groups, and should therefore be interpreted with some caution.

This global experiment revealed that nutrient enrichment, particularly nitrogen addition, can substantially reduce C_4_ dominance, including in regions where climatic conditions and initial community composition favour C_4_ plants. These results demonstrate that eutrophication and reduced herbivory can interact with the climatic thresholds underpinning C_4_ success in ways that are likely to reshape the structure and function of grassland ecosystems. While warming alone would be expected to favour C_4_ plants, concurrent increases in nutrient loading and changes in herbivory are likely to reverse this trajectory, reinforcing a potential global shift toward C_3_-dominated vegetation previously linked to increasing atmospheric CO_2_. Because C_4_ grasses sustain much of the planet’s primary productivity and grazing systems, their decline could reverberate through the global carbon cycle, food webs and food systems. Limiting nutrient pollution and protecting wild herbivore assemblages are thus crucial to safeguarding the world’s grasslands under a changing climate.

## Methods

### Site selection and experimental treatments

We tested our hypotheses using the Nutrient Network (www.nutnet.org), a globally replicated experiment quantifying the effects of herbivore exclusion and nutrient addition on grassland dynamics. To examine the role of climate (H1), we included data from 3,186 plots across 112 sites in the Nutrient Network^39^ (Fig. 5). The observational dataset (H1) represents a single sampling event at each site, with different sites sampled in different years (2007-2021; Supplementary Fig. 2). At all sites, photosynthetically active radiation (PAR) was measured with ceptometers above the canopy and at ground level. We calculated the proportion of PAR reaching the ground as a measure of the plot-level canopy openness. Because sites were sampled in different years, we tested the sensitivity of the best-supported model to sampling year by (i) including sampling year as a covariate and (ii) refitting the model to the 44 sites sampled in 2007-2008 (Supplementary Table 4).

**Figure 5.**
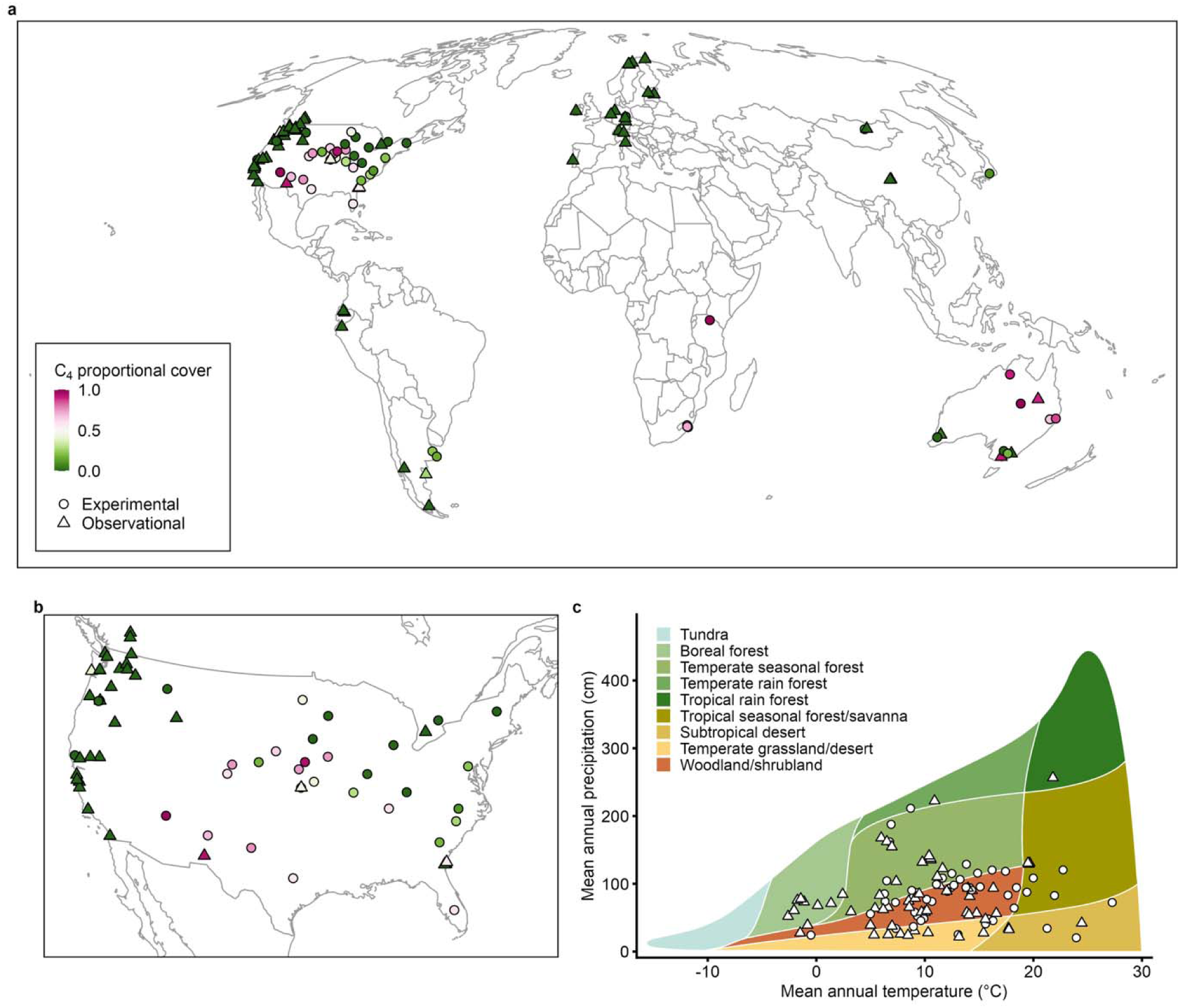
Distribution of study sites. a, Observational (triangles) and experimental (circles) sites, coloured by site-mean proportional C_4_ cover. b, Expanded view of North America, where the high density of sites makes individual locations difficult to distinguish in the global map. c, Sites in Whittaker biome space. European sites are not shown in detail because all sites there were C_3_-dominated. Projections: Mollweide (a) and Albers equal-area, NAD83 (b).

To test the nutrient and herbivory hypotheses (H2a-d), we used a subset of 48 experimental sites that received annual nutrient addition (N, P, K, and micronutrients (□)) and fencing treatments that exclude herbivores greater than approximately 50 g^39^. Fences do not exclude invertebrates or mammals smaller than approximately 50 g. Because the density and composition of excluded herbivores varied among sites, this treatment represents the removal of the locally present vertebrate herbivore assemblage. Analyses were restricted to sites that experience at least two months with climatic conditions meeting the crossover threshold defined in H1. We applied macronutrients at a rate of 10 g m^-2^ yr^-1^, together with micronutrients, which were only added in the first year. We refer to this combination as NPK throughout. We replicated each treatment across three to five blocks of ten 25 m^2^ plots per site. We included sites with at least one year of post-treatment data (median 6.5 years, maximum 15; Supplementary Table 7). In addition, 27 sites received a fully factorial combination of N, P, and K treatments, which allowed us to examine the effects of individual nutrients.

We surveyed plant composition annually in a 1 × 1 m permanent subplot within each 25 m² plot and visually estimated the percentage cover of individual species. To quantify C_4_ dominance, we calculated the proportional cover of C_4_ plants as the summed cover of all C_4_ species divided by combined cover of C_3_ and C_4_ species. To assign photosynthetic pathways, we combined Nutrient Network data with photosynthetic pathway information from the TRY database^69^. For species without species-level pathway data, we assigned pathways at the genus level when at least 90% of species in that genus with known pathways shared the same pathway. Species belonging to families with no known C_4_ or CAM lineages, and in genera that are uniformly C_3_, were assigned as C_3_^5,70–72^. Species that could not be assigned using this criterion, including unassigned species in genera known to be variable in pathway (e.g. *Cyperus*, *Euphorbia*, *Panicum*, *Salsola*) were excluded, as were species classed as C_3_–C_4_ intermediates or CAM. Excluded taxa accounted for 1.2% of vascular plant cover (0.1% CAM and intermediate species; Supplementary Table 11). Pathways were assigned at the species level for 85.1% of vascular plant cover and at the genus or family level for 13.8%; genus-level assignments made up 0.4% of C_4_ cover. Results were robust to assigning all unassigned taxa as C_3_ as well as to excluding genus-level assignments (Supplementary Table 10).

### Data analysis

C_4_ proportional cover (summed cover of C_4_ species divided by the combined cover of C_3_ and C_4_ species) included many exact zeroes and ones (Supplementary Fig. 1). We therefore used ordered beta regression^73^ with a logit link, implemented in glmmTMB^74^, for both observational (H1) and experimental (H2a-d) models in R v. 4.6.1^75^.

We tested whether C_4_ dominance followed climatic crossover thresholds (H1) by fitting ordered beta mixed models with the proportion of C_4_ cover as the response variable. We first examined relationships with conventional climate variables derived from long-term WorldClim averages^76^, including mean annual temperature, maximum temperature of the warmest month, mean annual precipitation, precipitation seasonality (the coefficient of variation of monthly precipitation) and the log ratio of warm-season to cool-season rainfall^20^. We then explicitly tested for climatic crossover thresholds by counting the number of months at each site that met combined temperature and precipitation criteria^10,18^.

We compared 110 threshold models, combining monthly average maximum temperatures of 22–32 °C with monthly precipitation of 15, 20, 25, 30 and 35 mm. For each combination, we counted either the number of months that met both criteria (55 models) or the numbers of warm months and wet months separately (55 models). We chose maximum temperatures to reflect daytime photosynthetic conditions^18^, and included precipitation to exclude hot but very dry months unlikely to support C_4_ growth^10^. Each model contained the threshold count(s) and the proportion of PAR reaching the ground, with random intercepts for site and block within site. Conventional climate variables were not included in these models because the threshold counts are derived from monthly temperature and precipitation. We selected the best-supported threshold using the Akaike Information Criterion (AIC; Supplementary Table 1, Supplementary Fig. 3). The best-supported threshold required both criteria to be met in the same month (≥23 °C and ≥20 mm). A threshold with the same temperature criterion but a lower precipitation threshold (≥15 mm) was also similarly supported (ΔAIC = 2.3), whereas models counting warm and wet months separately were much less supported (ΔAIC ≥ 21.0; Supplementary Table 1).

We compared the selected threshold model with models containing conventional climate variables, both individually and in combination, to test whether the threshold explained C_4_ cover beyond these variables (Supplementary Table 6). Mean annual precipitation improved the model only marginally and did not alter the threshold effect, so it was not retained in the final model (Supplementary Table 3). Variance inflation factors were all below 2. We tested for residual spatial autocorrelation using Moran’s I on site-level simulated residuals (DHARMa^77^) and by refitting the model with an exponential spatial covariance structure among sites (Supplementary Table 3). At the 86 sites with data from both sources, we compared the threshold model with an equivalent model in which the threshold count was replaced by the satellite-derived C_4_ distribution of Luo et al.^2^, comparing their marginal R^2^ values (Supplementary Table 2).

To test the temporal effects of nutrient addition and herbivore exclusion (H2), we compared control (unfertilized) plots with those receiving nutrient addition (NPK), both with and without herbivore exclusion. Models included experiment duration as a continuous predictor, both alone and in interaction with treatments. We restricted this analysis to sites that experienced at least two months meeting the crossover threshold identified in H1, resulting in 48 sites. We tested the sensitivity of this site selection by comparing results with two alternative filtering criteria (Supplementary Table 23): (1) excluding sites where the proportion cover of C_4_ species was consistently 0 throughout the experiment; and (2) excluding sites where mean pre-treatment C_4_ proportional cover was below 0.05 or above 0.95. All three subsets produced qualitatively similar results. We retained the threshold-only subset because it included sites where C_4_ species may initially have been absent but could plausibly occur given the climatic conditions and the regional species pool, as indicated by our observational analysis. Nine sites had no C_4_ cover in any analysed plot year but were retained for the same reason.

For each site, we calculated initial C4 cover as the mean plot-level C_4_ proportional cover across all plots in the experimental design, using pre-treatment data. To examine whether temporal trends in control plots depended on initial C4 cover, we used the fitted interaction model to estimate temporal slopes for control plots at relatively high and low initial C_4_ cover (mean ± 1 standard deviation). We assessed the significance of these conditional slopes by examining whether the 95% confidence intervals of the annual odds ratio excluded 1 (Supplementary Table 8). We then tested whether temporal declines in control plots reflected directional changes through calendar time rather than simply when the experiment began. We refitted ordered beta models to control plots, replacing plot age with calendar year (mean-centred) and including interactions with initial C_4_ cover and, in a second model, with the climatic threshold count. Random intercepts for site and plot within site were included in both models (Supplementary Table 9).

To isolate the effects of individual nutrients, we analysed the full treatment combinations (Control, N, P, K, NP, NK, PK, NPK, NPK+Fence, Fence) at the sites using the individual-nutrient design. We examined changes in C_4_ proportional cover between the pre-treatment survey (year 0) and year 4, representing five survey years. We selected this endpoint to allow treatment effects to emerge while retaining an adequate sample size. As in the main experimental analysis, we included only sites that experienced at least two months meeting the climatic crossover threshold identified in H1. Of the 34 such sites surveyed in both year 0 and year 4, we excluded seven at which C_4_ plants were absent from all plots in both years, as no change in C_4_ dominance was possible there. All plots with pre-treatment data were retained at the remaining 27 sites (836 plots), including plots without C_4_ cover in either year. To test whether the effects of N, P, and K were additive, we also fitted a factorial model including N, P, and K (each present or absent) and their interactions (excluding fenced plots). This model included random intercepts for site and blocks within site. At the 24 factorial sites with pre-treatment soil data, we tested whether treatment effects on C4 proportional cover depended on soil phosphorus (log-transformed ppm) or soil nitrogen (%), using separate linear mixed models with treatment × soil interactions and random intercepts for site and block within site (Supplementary Table 17). Pairwise differences among all ten treatments were estimated from the individual-nutrient model with Tukey adjustment (Supplementary Table 16).

Because proportional C_4_ cover does not distinguish between a decline in C_4_ cover and a compensatory increase in C_3_ cover, we also analysed the summed cover of C_4_ and C_3_ species separately. As summed cover can exceed 100% in multi-layered vegetation, we modelled absolute C_3_ and C_4_ cover using a Tweedie distribution and log link, following the same model structure as the proportional-cover models detailed below.

The observational models (H1) included random intercepts for site and for block within site, with continuous predictors standardised to a mean of 0 and standard deviation of 1. The experimental models (H2) included random intercepts for site, block within site and plot within block, to account for repeated measurements. Pre-treatment data were excluded from the experimental model response and used only to calculate initial C_4_ cover. Plot age was mean-centred, and initial C_4_ cover was mean-centred and scaled by its standard deviation. Functional-group proportions were modelled using the same approach as C4 proportional cover. Absolute C3 and C4 cover were modelled using Tweedie distributions with log link, while the proportion of PAR reaching the ground was modelled using beta regression (logit link) after transformation to the open interval^78^. The individual-nutrient and lineage change analyses used linear mixed models with random intercepts for site and block within site. We checked model residuals using DHARMa^77^. As a sensitivity analysis, we also fitted the main observational and experimental models using beta regression on transformed proportions (residual diagnostics in Supplementary Figs 4 and 6).

To explore how other plant groups responded to declining C_4_ species, we conducted two exploratory, post-hoc analyses. First, we modelled temporal changes in the proportional cover of C_3_ grasses, C_3_ forbs, and woody species across all treatment years using ordered beta mixed models, with treatment × year interaction and random intercepts for site, block within site and plot within block. Second, we examined whether treatment responses differed among major C_4_ grass lineages, which vary in life history and biogeographic distribution^64^. We separately modelled the treatment effects for Andropogoneae, Chloridoideae, the MPC clade (Melinidinae, Panicinae and Cenchrinae), and the genus *Aristida* - the four lineages with sufficient sample size to model individually (Supplementary Table 19). Although *Aristida* is a single genus, it represents a distinct lineage of C_4_ pathway evolution^71^. Additional independent C_4_ origins occur in grasses but were too sparsely represented to analyse^71^. Because the presence of functional groups and lineages varied among sites, each model was fitted to a different subset of sites. We report sample sizes with all estimates and interpret these results as exploratory.

The functional-group models were ordered beta mixed models of each group’s proportional cover across all treatment years, with a treatment × year interaction and random intercepts for site, block within site and plot within block. The individual lineage models were linear mixed models of changes in each lineage’s proportional cover between the pre-treatment year and year four, with random intercepts for site and block within site. To formally test whether responses differed among lineages, we combined data for the four lineages at sites where at least two co-occurred and compared linear mixed models of the change in proportional cover between the pre-treatment survey year and year four, with and without a lineage × treatment interaction. These models included random intercepts for site, block within site, and plot within block (Supplementary Table 22). We used a likelihood-ratio test to compare the models. Because these analyses were post hoc and involved multiple contrasts, we controlled the false-discovery rate across treatment contrasts within each set of models using the Benjamini-Hochberg procedure and report both raw and adjusted p-values.

## Data availability

Plot-level C_4_ proportional cover and plot-level cover summaries for functional groups, C_4_ grass lineage data, site-level climate and soil covariates, and all other associated components forming the minimum dataset to reproduce the analyses, figures and tables are available in figshare (https://figshare.com/s/4e524a1ec97d100e3477). Species-level cover data were contributed by Nutrient Network site investigators and are governed by the Nutrient Network data-use policy; they are available from the Nutrient Network coordinators on request via https://nutnet.org/data. Climate data were obtained from WorldClim v2.1 (https://www.worldclim.org), photosynthetic-pathway data from the TRY database (https://www.try-db.org) and C_4_ distribution data from Luo et al.^2^ All data underlying the figures and tables are provided in the figshare repository so no additional source data file is provided.

## Code availability

R code used to produce the analyses, figures, and tables is available in figshare at https://figshare.com/s/4e524a1ec97d100e3477

## Acknowledgements

This work was conducted using data from the Nutrient Network (http://www.nutnet.org) experiment, funded at the site scale by individual researchers. We thank the Minnesota Supercomputer Institute for hosting project data.

J.-C.S. and R.B. disclose funding from the Danish National Research Foundation (Center for Ecological Dynamics in a Novel Biosphere, ECONOVO; DNRF173). N.G.S. and E.E. disclose funding from the US National Science Foundation (DEB-2045968) and Texas Tech University. E.T.B. and E.W.S. disclose funding for Nutrient Network coordination and data management from the US National Science Foundation (DEB-1042132, DEB-1234162, DEB-1831944, DEB-2425352) and the Institute on the Environment (DG-0001-13). Y.M.B. discloses funding from Research Ireland, Northern Ireland’s Department of Agriculture, Environment and Rural Affairs (DAERA) and UK Research and Innovation (UKRI) through the International Science Partnerships Fund (ISPF; 22/CC/11103) at the Co-Centre for Climate + Biodiversity + Water. N.E. discloses funding from the German Centre for Integrative Biodiversity Research (iDiv) Halle–Jena–Leipzig, funded by the German Research Foundation (DFG; FZT 118, 202548816), and from the DFG (Ei 862/29-1). M.P. discloses funding from the Estonian Research Council (PRG3127) and the Estonian Ministry of Education and Research (Centre of Excellence AgroCropFuture, TK200). S.C.P. discloses funding from the US National Science Foundation through the Georgia Coastal Ecosystems Long-Term Ecological Research program (OCE-1832178). D.M.G. discloses funding from the US National Science Foundation (2550974). J.-C.S. also discloses funding from VILLUM Fonden (VILLUM Investigator project Biodiversity Dynamics in a Changing World; 16549) and the Independent Research Fund Denmark | Natural Sciences (MegaComplexity project; 0135-00225B). This is KBS contribution #X.

## Author contributions

J.A. and J.N.P. conceptualised the study, developed the methodology, analysed data, validated the analyses, collected data, wrote the first draft, edited drafts, created figures and contributed to project administration. R.B. and J.-C.S. developed the methodology, provided resources, wrote the first draft, edited drafts, supervised the work, contributed to project administration and provided funding. D.M.G. conceptualised the study, developed the methodology, analysed data, collected data, wrote the first draft, edited drafts and contributed to project administration. E.E. developed the methodology, analysed data and edited drafts. N.G.S. developed the methodology, collected data and edited drafts. E.T.B., Y.M.B., C.C., J.P.M., R.L.M., A.C.R., E.W.S., M.T. and G.M.W. collected data, provided resources, edited drafts and contributed to project administration. C.R.D., M.K., S.A.P. and S.M.P. collected data, edited drafts and contributed to project administration. L.L., P.M., H.M.M. and Z.R. collected data, provided resources and edited drafts. E.F.W. validated the analyses, collected data and edited drafts. C.B., L.A.B., M.N.B., M.C.C., S.C., I.D., N.E., K.J.E., A.E., M.G., S. Haider, N.H., S. Harpole, Y.H., A.J., J.M.H.K., S.E.K., K.J.K., A.D.B.L., M.M., A.S.M., J.W.M., M.P., S.C.P., P.L.P., C.R., M.S., R.S., R.V. and G.W. collected data and edited drafts.

## References

1. MacDougall, A. S. et al. The global extent of the grassland biome and implications for the terrestrial carbon sink. Nat Ecol Evol 10, 246–257 (2026).

2. Luo, X. et al. Mapping the global distribution of C4 vegetation using observations and optimality theory. Nat Commun 15, 1219 (2024).

3. Still, C. J., Berry, J. A., Collatz, G. J. & DeFries, R. S. Global distribution of C3 and C4 vegetation: Carbon cycle implications. Global Biogeochemical Cycles 17, 6-1-6–14 (2003).

4. Atkinson, R. R. L. et al. C4 photosynthesis boosts growth by altering physiology, allocation and size. Nature Plants 2, 16038 (2016).

5. Sage, R. F. A portrait of the C4 photosynthetic family on the 50th anniversary of its discovery: species number, evolutionary lineages, and Hall of Fame. Journal of Experimental Botany 68, e11–e28 (2017).

6. Barbehenn, R. V., Chen, Z., Karowe, D. N. & Spickard, A. C3 grasses have higher nutritional quality than C4 grasses under ambient and elevated atmospheric CO2. Global Change Biology 10, 1565–1575 (2004).

7. Nerlekar, A. N., Sullivan, L. L. & Brudvig, L. A. Grassland restorations must better foster forbs to facilitate high biodiversity. Restoration Ecology 32, e14214 (2024).

8. Piipponen, J. et al. Global trends in grassland carrying capacity and relative stocking density of livestock. Global Change Biology 28, 3902–3919 (2022).

9. Bremond, L., Boom, A. & Favier, C. Neotropical 3/4 grass distributions – present, past and future. Global Change Biology 18, 2324–2334 (2012).

10. Collatz, G. J., Berry, J. A. & Clark, J. S. Effects of climate and atmospheric CO2 partial pressure on the global distribution of C4 grasses: present, past, and future. Oecologia 114, 441–454 (1998).

11. Edwards, E. J. et al. The Origins of C4 Grasslands: Integrating Evolutionary and Ecosystem Science. Science 328, 587–591 (2010).

12. Ehleringer, J. R. Implications of quantum yield differences on the distributions of C3 and C4 grasses. Oecologia 31, 255–267 (1978).

13. Stowe, L. G. & Teeri, J. A. The Geographic Distribution of C4 Species of the Dicotyledonae in Relation to Climate. The American Naturalist https://doi.org/10.1086/283301 (1978) doi:10.1086/283301.

14. Young, S. N. R., Dunning, L. T., Liu, H., Stevens, C. J. & Lundgren, M. R. C4 trees have a broader niche than their close C3 relatives. J Exp Bot 73, 3189–3204 (2022).

15. Bond, W. J. Large parts of the world are brown or black: A different view on the ‘Green World’hypothesis. Journal of Vegetation Science 16, 261–266 (2005).

16. Lehmann, C. E. R., Archibald, S. A., Hoffmann, W. A. & Bond, W. J. Deciphering the distribution of the savanna biome. New Phytologist 191, 197–209 (2011).

17. Monson, R. K. et al. C4 photosynthesis, trait spectra, and the fast-efficient phenotype. New Phytologist 246, 879–893 (2025).

18. Griffith, D. M. et al. Biogeographically distinct controls on C3 and C4 grass distributions: merging community and physiological ecology. Global Ecology and Biogeography 24, 304– 313 (2015).

19. Hattersley, P. W. The distribution of C3 and C4 grasses in Australia in relation to climate. Oecologia 57, 113–128 (1983).

20. Xie, Q. et al. Satellite-observed shifts in C3/C4 abundance in Australian grasslands are associated with rainfall patterns. Remote Sensing of Environment 273, 112983 (2022).

21. Brown, R. H. A Difference in N Use Efficiency in C3 and C4 Plants and its Implications in Adaptation and Evolution. Crop Science 18, cropsci1978.0011183X001800010025x (1978).

22. Long, S. P. Environmental responses. C4 plant biology 215–249 (1999).

23. Taylor, S. H. et al. Ecophysiological traits in C3 and C4 grasses: a phylogenetically controlled screening experiment. New Phytologist 185, 780–791 (2010).

24. Tieszen, L., Senyimba, M., Imbamba, S. & Troughton, J. The distribution of C3 and C4 grasses and carbon isotope discrimination along an altitudinal and moisture gradient in Kenya. Oecologia 37, 337–350 (1979).

25. Waring, E. F., Perkowski, E. A. & Smith, N. G. Soil nitrogen fertilization reduces relative leaf nitrogen allocation to photosynthesis. J Exp Bot 74, 5166–5180 (2023).

26. Stevens, N., Bond, W., Feurdean, A. & Lehmann, C. E. R. Grassy Ecosystems in the Anthropocene. Annual Review of Environment and Resources 47, 261–289 (2022).

27. Stevens, C. J., David, T. I. & Storkey, J. Atmospheric nitrogen deposition in terrestrial ecosystems: Its impact on plant communities and consequences across trophic levels. Functional Ecology 32, 1757–1769 (2018).

28. Harpole, W. S. et al. Nutrient co-limitation of primary producer communities. Ecology Letters 14, 852–862 (2011).

29. Fay, P. A. et al. Grassland productivity limited by multiple nutrients. Nature Plants 1, 15080 (2015).

30. Griffith, D. M., Cotton, J. M., Powell, R. L., Sheldon, N. D. & Still, C. J. Multi-century stasis in C3 and C4 grass distributions across the contiguous United States since the industrial revolution. Journal of Biogeography 44, 2564–2574 (2017).

31. Oberhuber, W., Dai, Z.-Y. & Edwards, G. E. Light dependence of quantum yields of Photosystem II and CO2 fixation in C3 and C4 plants. Photosynth Res 35, 265–274 (1993).

32. Skillman, J. B. Quantum yield variation across the three pathways of photosynthesis: not yet out of the dark. J Exp Bot 59, 1647–1661 (2008).

33. Hoetzel, S., Dupont, L., Schefuß, E., Rommerskirchen, F. & Wefer, G. The role of fire in Miocene to Pliocene C4 grassland and ecosystem evolution. Nature Geosci 6, 1027–1030 (2013).

34. Karp, A. T., Behrensmeyer, A. K. & Freeman, K. H. Grassland fire ecology has roots in the late Miocene. Proceedings of the National Academy of Sciences 115, 12130–12135 (2018).

35. Keeley, J. E. & Rundel, P. W. Fire and the Miocene expansion of C4 grasslands. Ecology Letters 8, 683–690 (2005).

36. Fanselow, N. et al. Short-term regrowth responses of four steppe grassland species to grazing intensity, water and nitrogen in Inner Mongolia. Plant Soil 340, 279–289 (2011).

37. Zheng, S. et al. Differential responses of plant functional trait to grazing between two contrasting dominant C3 and C4 species in a typical steppe of Inner Mongolia, China. Plant Soil 340, 141–155 (2011).

38. Augustine, D. J., Derner, J. D., Milchunas, D., Blumenthal, D. & Porensky, L. M. Grazing moderates increases in C3 grass abundance over seven decades across a soil texture gradient in shortgrass steppe. Journal of Vegetation Science 28, 562–572 (2017).

39. Borer, E. T. et al. Finding generality in ecology: a model for globally distributed experiments. Methods in Ecology and Evolution 5, 65–73 (2014).

40. Knapp, A. K. et al. Resolving the Dust Bowl paradox of grassland responses to extreme drought. Proceedings of the National Academy of Sciences 117, 22249–22255 (2020).

41. Simpson, K. J. et al. Increasing CO2 levels fertilize C4 grass production. Nature 658, 415– 420 (2026).

42. Reich, P. B., Hobbie, S. E., Lee, T. D. & Pastore, M. A. Unexpected reversal of C3 versus C4 grass response to elevated CO2 during a 20-year field experiment. Science 360, 317– 320 (2018).

43. Borer, E. T. & Stevens, C. J. Nitrogen deposition and climate: an integrated synthesis. Trends in Ecology & Evolution 37, 541–552 (2022).

44. Simkin, S. M. et al. Conditional vulnerability of plant diversity to atmospheric nitrogen deposition across the United States. Proceedings of the National Academy of Sciences 113, 4086–4091 (2016).

45. Tilman, D. Nitrogen-Limited Growth in Plants from Different Successional Stages. Ecology 67, 555–563 (1986).

46. Wedin, D. A. & Tilman, D. Influence of Nitrogen Loading and Species Composition on the Carbon Balance of Grasslands. Science 274, 1720–1723 (1996).

47. Reay, D. S., Dentener, F., Smith, P., Grace, J. & Feely, R. A. Global nitrogen deposition and carbon sinks. Nature Geosci 1, 430–437 (2008).

48. Midolo, G. et al. Sixty years of plant community change in Europe indicate a shift toward nutrient-richer and denser vegetation. Science Advances 12, eaeb2493 (2026).

49. Trepel, J. et al. Global analysis suggests nitrogen deposition as an underestimated driver of vegetation greening. Ecography https://doi.org/10.1002/ecog.08631 (2026) doi:10.1002/ecog.08631.

50. Zhu, J. et al. Changing patterns of global nitrogen deposition driven by socio-economic development. Nat Commun 16, 46 (2025).

51. Vitousek, P. M., Porder, S., Houlton, B. Z. & Chadwick, O. A. Terrestrial phosphorus limitation: mechanisms, implications, and nitrogen–phosphorus interactions. Ecological Applications 20, 5–15 (2010).

52. Borer, E. T. et al. Herbivores and nutrients control grassland plant diversity via light limitation. Nature 508, 517–520 (2014).

53. Borer, E. T. et al. Nutrients cause grassland biomass to outpace herbivory. Nat Commun 11, 6036 (2020).

54. Fløjgaard, C., Pedersen, P. B. M., Sandom, C. J., Svenning, J.-C. & Ejrnæs, R. Exploring a natural baseline for large-herbivore biomass in ecological restoration. Journal of Applied Ecology 59, 18–24 (2022).

55. Zhang, Q. et al. Grazing primarily drives the relative abundance change of C4 plants in the typical steppe grasslands across households at a regional scale. Rangel. J. 36, 565–572 (2014).

56. Derner, J. D., Boutton, T. W. & Briske, D. D. Grazing and Ecosystem Carbon Storage in the North American Great Plains. Plant Soil 280, 77–90 (2006).

57. Auerswald, K. et al. C4 abundance in an Inner Mongolia grassland system is driven by temperature–moisture interaction, not grazing pressure. Basic and Applied Ecology 13, 67– 75 (2012).

58. Blumenthal, D. M. et al. Traits link drought resistance with herbivore defence and plant economics in semi-arid grasslands: The central roles of phenology and leaf dry matter content. Journal of Ecology 108, 2336–2351 (2020).

59. Price, J. N. et al. Evolutionary history of grazing and resources determine herbivore exclusion effects on plant diversity. Nat Ecol Evol 6, 1290–1298 (2022).

60. Koerner, S. E. et al. Change in dominance determines herbivore effects on plant biodiversity. Nat Ecol Evol 2, 1925–1932 (2018).

61. Nelson, R. A. et al. Forb diversity globally is harmed by nutrient enrichment but can be rescued by large mammalian herbivory. Commun Biol 8, 444 (2025).

62. Furey, G. N., Hawthorne, P. L. & Tilman, D. Might field experiments also be inadvertent metacommunities? Ecology 103, e3694 (2022).

63. Griffith, D. M. et al. Lineage-based functional types: characterising functional diversity to enhance the representation of ecological behaviour in Land Surface Models. New Phytologist 228, 15–23 (2020).

64. Lehmann, C. E. R. et al. Functional diversification enabled grassy biomes to fill global climate space. 583625 Preprint at 10.1101/583625 (2019).

65. Craine, J. M. et al. Functional traits, productivity and effects on nitrogen cycling of 33 grassland species. Functional Ecology 16, 563–574 (2002).

66. Hsu, S.-K. et al. Contrasting rhizosphere nitrogen dynamics in Andropogoneae grasses. The Plant Journal 123, e70319 (2025).

67. Bakker, E. S., Ritchie, M. E., Olff, H., Milchunas, D. G. & Knops, J. M. H. Herbivore impact on grassland plant diversity depends on habitat productivity and herbivore size. Ecology Letters 9, 780–788 (2006).

68. Olff, H. & Ritchie, M. E. Effects of herbivores on grassland plant diversity. Trends in Ecology & Evolution 13, 261–265 (1998).

69. Kattge, J. et al. TRY Plant Trait Database – Enhanced Coverage and Open Access. Global Change Biology 26, 119–188 (2020).

70. Sage, R. F., Christin, P.-A. & Edwards, E. J. The C4 plant lineages of planet Earth. Journal of Experimental Botany 62, 3155–3169 (2011).

71. Grass Phylogeny Working Group. New grass phylogeny resolves deep evolutionary relationships and discovers C4 origins. New Phytologist 193, 304–312 (2012).

72. Silvera, K. et al. Evolution along the crassulacean acid metabolism continuum. Functional Plant Biology 37, 995–1010 (2010).

73. Kubinec, R. Ordered Beta Regression: A Parsimonious, Well-Fitting Model for Continuous Data with Lower and Upper Bounds. Political Analysis 31, 519–536 (2023).

74. Brooks, M. et al. glmmTMB: Generalized Linear Mixed Models using Template Model Builder. (2024).

75. R Core Team. R: A Language and Environment for Statistical Computing. (R Foundation for Statistical Computing, Vienna, Austria, 2024).

76. Fick, S. E. & Hijmans, R. J. WorldClim 2: New 1-Km Spatial Resolution Climate Surfaces for Global Land Areas. International Journal of Climatology 37, 4302–4315 (2017).

77. Hartig, F., Lohse, L., Leite, M. de S. & Werneke, C. DHARMa: Residual Diagnostics for Hierarchical (Multi-Level / Mixed) Regression Models. (2026).

78. Smithson, M. & Verkuilen, J. A better lemon squeezer? Maximum-likelihood regression with beta-distributed dependent variables. Psychol Methods 11, 54–71 (2006).

