## Supplementary Information for "Nutrient enrichment and herbivore exclusion disrupt the climate-driven balance between C_3_ and C_4_ plants"

Joe Atkinson[1,2,*,†], Jodi N Price[3,†], Robert Buitenwerf[2], Nicholas G Smith[4], Ezinwanne Ezekannagha[4], Elizabeth T Borer[5], Cynthia Brown[6], Lars A Brudvig[7], Yvonne M Buckley[8], Miguel N Bugalho[9], Maria C Caldeira[10], Sofía Campana[11], Clinton Carbutt[12,13], Chris R Dickman[14], Ian Donohue[15], Nico Eisenhauer[16,17], Kenneth J Elgersma[18], Anu Eskelinen[16,19], Magda Garbowski[20], Sylvia Haider[21], Nicole Hagenah[22], Stanley Harpole[16,23,24], Yann Hautier[25], Anke Jentsch[26], Johannes MH Knops[27], Sally E Koerner[28], Mayank Kohli[29], Kimberly J Komatsu[28], Lauri Laanisto[30], Andrew DB Leakey[31], Petr Macek[30,32], Miaojun Ma[33,34], Andrew S MacDougall[35], Jason P Martina[36], Holly M Martinson[37], Rebecca L McCulley[38], John W Morgan[39], Meelis Pärtel[40], Steven C Pennings[41], Pablo L Peri[42], Sally A Power[43], Suzanne M Prober[44], Zhengwei Ren[33,34], Anita C Risch[45], Christiane Roscher[16,46], Mahesh Sankaran[29], Eric W Seabloom[5], Rachel Standish[47], Michelle Tedder[12], Risto Virtanen[19], Glenda M Wardle[14,48], Elizabeth F Waring[49], George Wheeler[27,50], Daniel M Griffith[51], Jens-Christian Svenning[2]

†These authors contributed equally: Joe Atkinson, Jodi N Price

1. School of Biological Sciences, Adelaide University, Adelaide, South Australia, Australia
2. Center for Ecological Dynamics in a Novel Biosphere (ECONOVO), Department of Biology, Aarhus University, Aarhus, Denmark
3. Gulbali Institute, Charles Sturt University, Albury, New South Wales, Australia
4. Department of Biological Sciences, Texas Tech University, Lubbock, TX USA
5. Department of Ecology, Evolution, and Behavior, University of Minnesota, St. Paul, MN 55108 USA
6. Department of Agricultural Biology, Colorado State University, Fort Collins, CO, USA
7. Department of Plant Biology and Program in Ecology, Evolution, and Behavior, Michigan State University, East Lansing, MI 48824, USA
8. Co-Centre for Climate + Biodiversity + Water, Trinity College Dublin, Dublin 2, Ireland
9. Centre for Applied Ecology ”Prof. Baeta Neves” (CEABN-InBIO), School of Agriculture, University of Lisbon, Portugal
10. Forest Research Centre, Associate Laboratory TERRA, School of Agriculture, University of Lisbon, Portugal
11. Universidad de Buenos Aires, Facultad de Agronomía, Departamento de Recursos Naturales y Ambiente, Cátedra de Ecología. CONICET. Instituto de Investigaciones Fisiológicas y Ecológicas Vinculadas a la Agricultura (IFEVA). Buenos Aires, Argentina
12. School of Agriculture and Science, University of KwaZulu-Natal, Scottsville 3209, South Africa
13. Scientific Services, Ezemvelo KZN Wildlife, Cascades 3202, South Africa
14. School of Life and Environmental Sciences, The University of Sydney, NSW 2006, Australia
15. School of Natural Sciences, Trinity College Dublin, Dublin, Ireland
16. German Centre for Integrative Biodiversity Research (iDiv) Halle-Jena-Leipzig, Puschstrasse 4, 04103 Leipzig, Germany
17. Institute of Biology, Leipzig University, Puschstrasse 4, 04103 Leipzig, Germany
18. Department of Biology, University of Northern Iowa, Cedar Falls, IA, USA
19. Ecology and Genetics Unit, University of Oulu, Finland
20. Department of Animal and Range Sciences, New Mexico State University, Las Cruces, NM, USA
21. Institute of Ecology, School of Sustainability, Leuphana University of Lüneburg, Germany
22. Mammal Research Institute, Department of Zoology and Entomology, University of Pretoria, Pretoria, South Africa
23. Helmholtz Centre for Environmental Research -UFZ, Department Physiological Diversity, Permoserstrasse 14, 04318 Leipzig, Germany
24. Martin Luther University Halle-Wittenberg, am Kirchtor 1, 06108 Halle (Saale), Germany

25. Ecology and Biodiversity Group, Department of Biology, Utrecht University, Utrecht, Netherlands
26. Department of Disturbance Ecology and Vegetation Dynamics, University of Bayreuth, Germany
27. School of Biological Sciences, University of Nebraska-Lincoln, Lincoln, NE 68588, USA
28. Department of Biology, University of North Carolina at Greensboro, Greensboro, NC 27402, USA
29. National Centre for Biological Sciences, TIFR, Bellary Road, Bengaluru 560065, Karnataka, India
30. Chair of Biodiversity and Nature Tourism, Estonian University of Life Sciences, Tartu, Estonia
31. Department of Plant Biology, University of Illinois Urbana-Champaign, Urbana IL, USA
32. Institute of Hydrobiology, Biology Centre of the Czech Academy of Sciences, Na Sadkach 7, Ceske Budejovice, Czech Republic
33. College of Ecology, Lanzhou University, No. 222 Tianshui South Road, Lanzhou City, China
34. The Gansu Gannan Grassland Ecosystem National Observation and Research Station, Maqu County, Gansu Province, China
35. Department of Integrative Biology, University of Guelph, Guelph, ON, N1G2W1, Canada
36. Department of Biology, Texas State University, San Marcos, TX 78666, USA
37. Department of Biology, McDaniel College, Westminster, MD, USA
38. Department of Plant & Soil Sciences, University of Kentucky, Lexington, KY, 40546-0312, USA
39. Department of Environment and Genetics, La Trobe University, Bundoora, Victoria, Australia
40. Institute of Ecology and Earth Sciences, University of Tartu, Tartu, Estonia
41. Department of Biology and Biochemistry, University of Houston, Houston TX, 77204, USA
42. Universidad Nacional de la Patagonia Austral-Instituto Nacional de Tecnología, Agropecuaria-Consejo Nacional de Investigaciones Científicas y Técnicas, Rio Gallegos CP, 9400, Santa Cruz, Argentina
43. Hawkesbury Institute for the Environment, Western Sydney University, Locked Bag 1797, Penrith, New South Wales, 2751, Australia
44. CSIRO Environment, Canberra, ACT 2601 Australia
45. Swiss Federal Institute for Forest, Snow and Landscape Research WSL, 8903 Birmensdorf, Switzerland
46. Helmholtz Centre for Environmental Research -UFZ, Department Biodiversity and People, Permoserstrasse 14, 04318 Leipzig, Germany
47. School of Environmental and Conservation Sciences, Murdoch University, 90 South Street, Murdoch, Western Australia
48. ARC Training Centre in Data Analytics for Resources and Environments (DARE), The University of Sydney, NSW 2006, Australia
49. Department of Biological Sciences, Northeastern State University, Tahlequah, OK USA
50. Department of Biological Sciences, Michigan Technological University, Houghton, MI, USA
51. Department of Ecology and Evolution, Stony Brook University, Stony Brook, NY 11794, USA

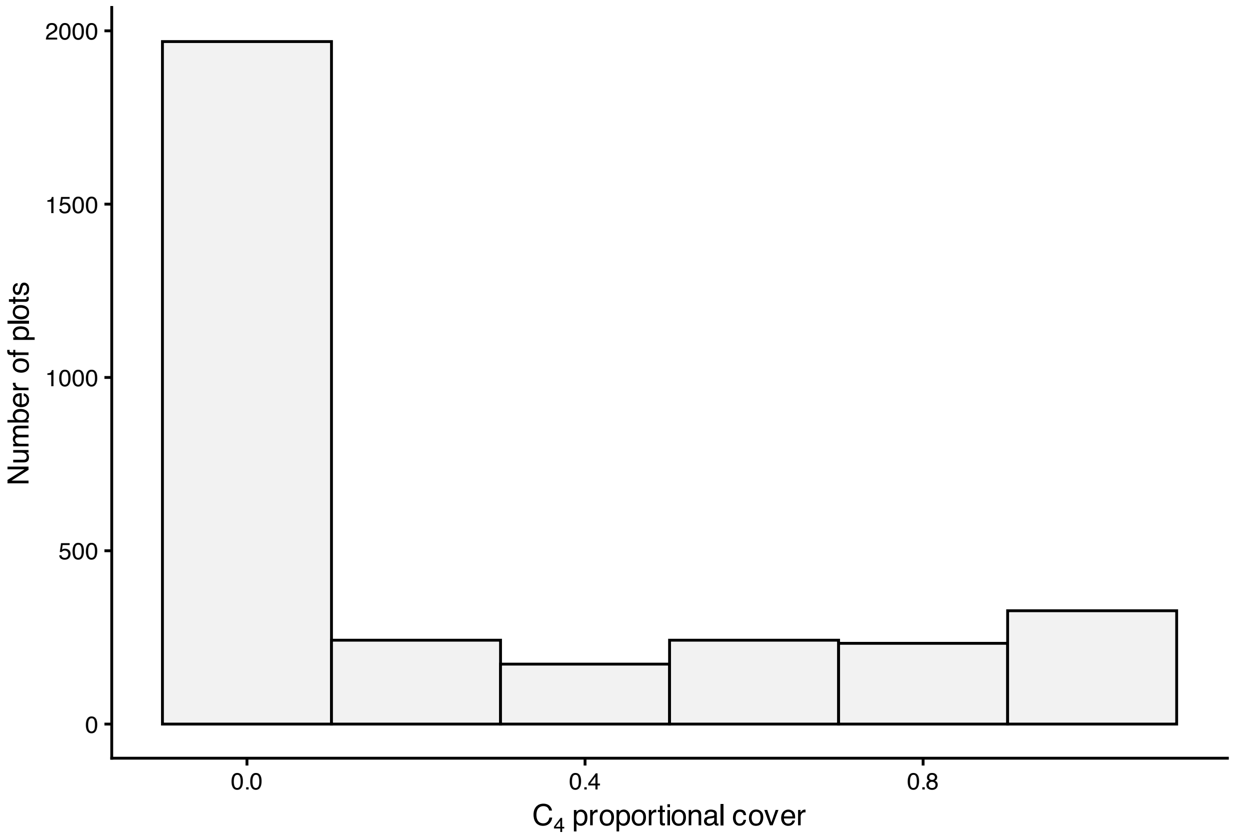


**Supplementary Fig. 1.** Distribution of the response variable $C_4$ proportional cover.

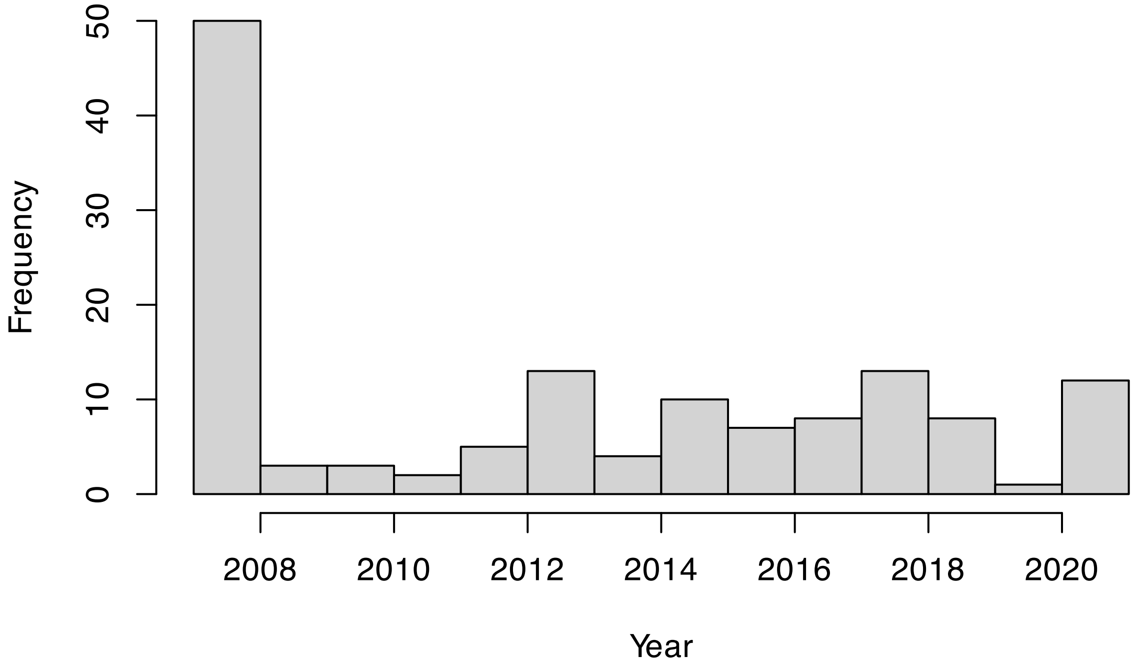


**Supplementary Fig. 2.** Distribution of survey years in the observational dataset.

Models test all combinations of temperature (22-32 °C) and precipitation (15-35 mm) thresholds to identify the best-fitting crossover threshold.

**Supplementary Table 1.** Model selection for the climatic crossover threshold. “Same month” models count months in which both the temperature and the precipitation criteria were met. “separate counts” models include a count of warm months and a count of wet months as two predictors. The baseline model has no threshold count. Models are ordered beta mixed models of $C_4$ proportional cover containing the threshold count and ground-level PAR, with site and block random intercepts. Conventional climate variables were not included because the counts are derived from temperature and precipitation (they are compared with the threshold in Supplementary Table 6).

| Type | Threshold | ΔAIC | Weight |
|---|---|---|---|
| Same month | >= 23 °C, >= 20 mm | 0.0 | 0.434 |
| Same month | >= 23 °C, >= 15 mm | 2.3 | 0.136 |
| Same month | >= 23 °C, >= 25 mm | 3.7 | 0.067 |
| Same month | >= 24 °C, >= 20 mm | 4.4 | 0.047 |
| Same month | >= 26 °C, >= 20 mm | 4.6 | 0.043 |
| Same month | >= 25 °C, >= 20 mm | 4.8 | 0.039 |
| Same month | >= 24 °C, >= 25 mm | 5.4 | 0.029 |
| Same month | >= 26 °C, >= 30 mm | 5.6 | 0.026 |
| Same month | >= 23 °C, >= 30 mm | 6.0 | 0.022 |
| Same month | >= 24 °C, >= 35 mm | 6.1 | 0.020 |
| Same month | >= 23 °C, >= 35 mm | 6.2 | 0.019 |
| Same month | >= 26 °C, >= 15 mm | 6.3 | 0.019 |
| Same month | >= 24 °C, >= 30 mm | 6.5 | 0.017 |
| Same month | >= 26 °C, >= 35 mm | 7.0 | 0.013 |
| Same month | >= 25 °C, >= 35 mm | 7.1 | 0.012 |
| Same month | >= 26 °C, >= 25 mm | 7.1 | 0.012 |
| Same month | >= 25 °C, >= 30 mm | 7.6 | 0.010 |
| Same month | >= 24 °C, >= 15 mm | 8.2 | 0.007 |
| Same month | >= 25 °C, >= 25 mm | 8.3 | 0.007 |
| Same month | >= 25 °C, >= 15 mm | 8.5 | 0.006 |
| Same month | >= 22 °C, >= 20 mm | 8.8 | 0.005 |
| Same month | >= 22 °C, >= 25 mm | 9.2 | 0.004 |
| Same month | >= 22 °C, >= 15 mm | 10.4 | 0.002 |
| Same month | >= 22 °C, >= 35 mm | 11.9 | 0.001 |
| Same month | >= 22 °C, >= 30 mm | 11.9 | 0.001 |
| Same month | >= 27 °C, >= 20 mm | 18.6 | <0.001 |
| Same month | >= 27 °C, >= 30 mm | 19.0 | <0.001 |
| Same month | >= 27 °C, >= 15 mm | 19.5 | <0.001 |
| Same month | >= 27 °C, >= 25 mm | 19.9 | <0.001 |
| Same month | >= 27 °C, >= 35 mm | 21.0 | <0.001 |
| Separate | >= 23 °C, >= 35 mm | 21.0 | <0.001 |
| Separate | >= 23 °C, >= 30 mm | 21.4 | <0.001 |
| Separate | >= 23 °C, >= 25 mm | 21.6 | <0.001 |
| Separate | >= 23 °C, >= 20 mm | 21.7 | <0.001 |
| Separate | >= 23 °C, >= 15 mm | 21.7 | <0.001 |
| Same month | >= 28 °C, >= 20 mm | 23.0 | <0.001 |
| Same month | >= 28 °C, >= 15 mm | 23.3 | <0.001 |
| Separate | >= 24 °C, >= 35 mm | 24.2 | <0.001 |
| Separate | >= 24 °C, >= 30 mm | 24.5 | <0.001 |
| Separate | >= 24 °C, >= 15 mm | 24.5 | <0.001 |
| Separate | >= 24 °C, >= 25 mm | 24.6 | <0.001 |
| Separate | >= 24 °C, >= 20 mm | 24.6 | <0.001 |
| Same month | >= 28 °C, >= 30 mm | 24.7 | <0.001 |
| Same month | >= 28 °C, >= 25 mm | 25.0 | <0.001 |
| Same month | >= 28 °C, >= 35 mm | 25.5 | <0.001 |
| Separate | >= 22 °C, >= 35 mm | 27.0 | <0.001 |
| Same month | >= 29 °C, >= 20 mm | 27.3 | <0.001 |
| Separate | >= 22 °C, >= 30 mm | 27.5 | <0.001 |
| Separate | >= 22 °C, >= 25 mm | 27.7 | <0.001 |
| Separate | >= 22 °C, >= 15 mm | 27.7 | <0.001 |
| Separate | >= 22 °C, >= 20 mm | 27.8 | <0.001 |
| Same month | >= 29 °C, >= 15 mm | 27.8 | <0.001 |
| Same month | >= 29 °C, >= 30 mm | 28.2 | <0.001 |
| Same month | >= 29 °C, >= 35 mm | 28.7 | <0.001 |
| Same month | >= 29 °C, >= 25 mm | 29.7 | <0.001 |
| Separate | >= 26 °C, >= 35 mm | 30.1 | <0.001 |
| Separate | >= 25 °C, >= 35 mm | 30.3 | <0.001 |
| Separate | >= 26 °C, >= 30 mm | 30.4 | <0.001 |
| Separate | >= 26 °C, >= 25 mm | 30.5 | <0.001 |
| Separate | >= 25 °C, >= 30 mm | 30.6 | <0.001 |
| Separate | >= 26 °C, >= 20 mm | 30.6 | <0.001 |
| Separate | >= 26 °C, >= 15 mm | 30.7 | <0.001 |
| Separate | >= 25 °C, >= 15 mm | 30.7 | <0.001 |
| Separate | >= 25 °C, >= 25 mm | 30.7 | <0.001 |
| Separate | >= 25 °C, >= 20 mm | 30.7 | <0.001 |
| Same month | >= 30 °C, >= 30 mm | 35.7 | <0.001 |
| Same month | >= 30 °C, >= 35 mm | 35.7 | <0.001 |
| Same month | >= 30 °C, >= 15 mm | 36.7 | <0.001 |
| Same month | >= 30 °C, >= 20 mm | 36.9 | <0.001 |
| Separate | >= 27 °C, >= 35 mm | 37.6 | <0.001 |
| Separate | >= 27 °C, >= 30 mm | 38.0 | <0.001 |
| Separate | >= 27 °C, >= 25 mm | 38.1 | <0.001 |
| Same month | >= 30 °C, >= 25 mm | 38.1 | <0.001 |
| Separate | >= 27 °C, >= 20 mm | 38.1 | <0.001 |
| Separate | >= 27 °C, >= 15 mm | 38.2 | <0.001 |
| Separate | >= 28 °C, >= 35 mm | 42.6 | <0.001 |
| Separate | >= 28 °C, >= 30 mm | 42.9 | <0.001 |
| Separate | >= 28 °C, >= 25 mm | 42.9 | <0.001 |
| Separate | >= 28 °C, >= 20 mm | 43.0 | <0.001 |
| Separate | >= 28 °C, >= 15 mm | 43.0 | <0.001 |
| Same month | >= 31 °C, >= 35 mm | 46.9 | <0.001 |
| Same month | >= 31 °C, >= 30 mm | 47.5 | <0.001 |
| Separate | >= 29 °C, >= 35 mm | 47.7 | <0.001 |
| Separate | >= 29 °C, >= 30 mm | 48.0 | <0.001 |
| Separate | >= 29 °C, >= 25 mm | 48.1 | <0.001 |
| Separate | >= 29 °C, >= 20 mm | 48.2 | <0.001 |
| Separate | >= 29 °C, >= 15 mm | 48.3 | <0.001 |
| Same month | >= 31 °C, >= 15 mm | 48.4 | <0.001 |
| Same month | >= 31 °C, >= 20 mm | 48.4 | <0.001 |
| Same month | >= 31 °C, >= 25 mm | 49.6 | <0.001 |
| Separate | >= 30 °C, >= 35 mm | 49.6 | <0.001 |
| Separate | >= 30 °C, >= 30 mm | 49.8 | <0.001 |
| Separate | >= 30 °C, >= 20 mm | 49.8 | <0.001 |
| Separate | >= 30 °C, >= 25 mm | 49.8 | <0.001 |
| Separate | >= 30 °C, >= 15 mm | 49.9 | <0.001 |
| Same month | >= 32 °C, >= 35 mm | 55.6 | <0.001 |
| Separate | >= 31 °C, >= 35 mm | 55.9 | <0.001 |
| Separate | >= 31 °C, >= 30 mm | 55.9 | <0.001 |
| Separate | >= 31 °C, >= 20 mm | 56.0 | <0.001 |
| Separate | >= 31 °C, >= 15 mm | 56.0 | <0.001 |
| Separate | >= 31 °C, >= 25 mm | 56.0 | <0.001 |
| Same month | >= 32 °C, >= 30 mm | 56.2 | <0.001 |
| Same month | >= 32 °C, >= 15 mm | 56.4 | <0.001 |
| Same month | >= 32 °C, >= 20 mm | 57.4 | <0.001 |
| Same month | >= 32 °C, >= 25 mm | 58.1 | <0.001 |
| Separate | >= 32 °C, >= 35 mm | 61.5 | <0.001 |
| Separate | >= 32 °C, >= 30 mm | 61.5 | <0.001 |
| Separate | >= 32 °C, >= 15 mm | 61.6 | <0.001 |
| Separate | >= 32 °C, >= 20 mm | 61.6 | <0.001 |
| Separate | >= 32 °C, >= 25 mm | 61.6 | <0.001 |
| | No threshold term | 75.8 | <0.001 |

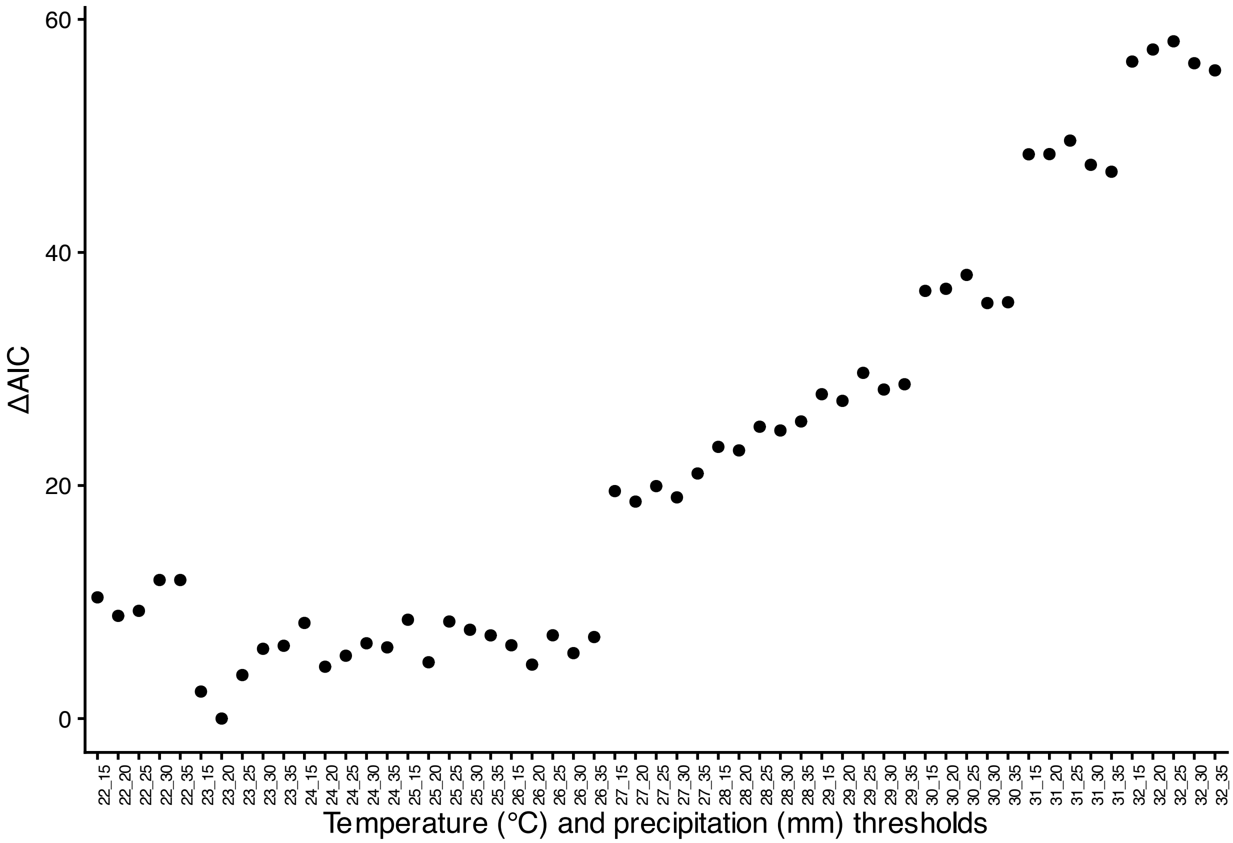


**Supplementary Fig. 3.** ΔAIC for the same-month temperature-precipitation crossover threshold models (ordered beta mixed models with the threshold count and ground-level PAR; Supplementary Table 1). Labels give the temperature (°C) and precipitation (mm) criteria.

**Supplementary Table 2.** Comparison of the crossover threshold model with the predicted $C_4$ area of Luo et al.[1].

| | p raw | | | | p raw | | | |
|---|---|---|---|---|---|---|---|---|
| *Predictors* | *Estimates* | *CI* | *Statistic* | *p* | *Estimates* | *CI* | *Statistic* | *p* |
| Intercept | 0.003 | 0.001 – 0.009 | -9.812 | <0.001 | 0.000 | 0.000 – 0.001 | -7.469 | <0.001 |
| Proportion of PAR reaching the ground | 0.990 | 0.880 – 1.113 | -0.175 | 0.861 | 0.984 | 0.875 – 1.108 | -0.263 | 0.792 |
| Months ≥ 23 °C and ≥ 20 mm | 95.659 | 34.156 – 267.904 | 8.680 | <0.001 | | | | |
| Luo et al. $C_4$ area | | | | | 1.231 | 1.141 – 1.327 | 5.381 | <0.001 |
| **Random Effects** | | | | | | | | |
| $\sigma^2$ | 3.58 | | | | 3.58 | | | |
| $\tau_{00}$ | 0.27 $_{\text{block:site\_code}}$ | | | | 0.27 $_{\text{block:site\_code}}$ | | | |
| | 15.23 $_{\text{site\_code}}$ | | | | 27.15 $_{\text{site\_code}}$ | | | |
| ICC | 0.81 | | | | 0.88 | | | |
| N | 6 $_{\text{block}}$ | | | | 6 $_{\text{block}}$ | | | |
| | 86 $_{\text{site\_code}}$ | | | | 86 $_{\text{site\_code}}$ | | | |
| Observations | 2548 | | | | 2548 | | | |
| Marginal $R^2$ / Conditional $R^2$ | 0.519 / 0.910 | | | | 0.267 / 0.915 | | | |

Based on Supplementary Table 1, the best-fitting model uses months where mean maximum temperature is

≥ 23 °C and mean precipitation is ≥ 20 mm in the same month.

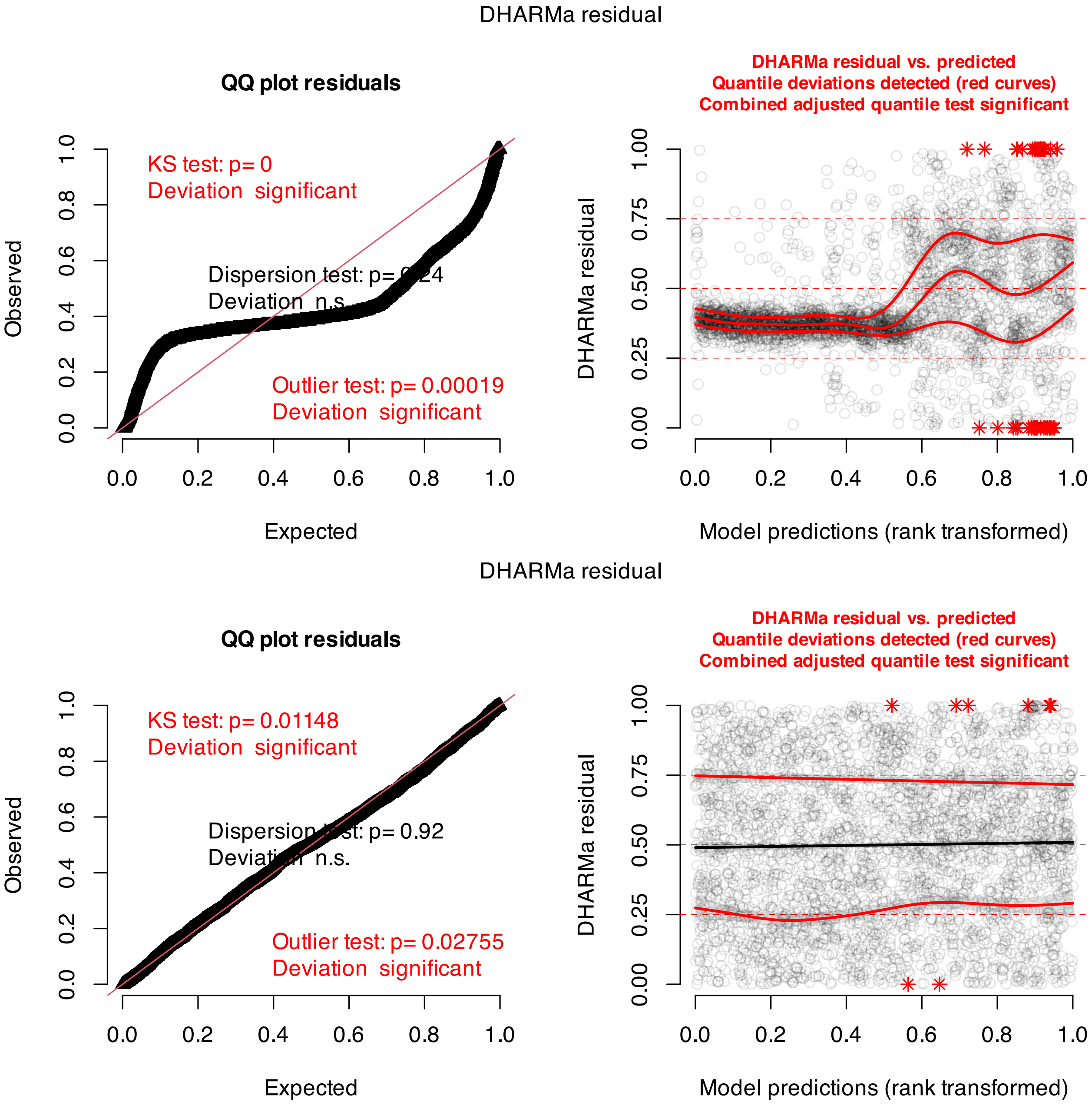


**Supplementary Fig. 4.** DHARMa[2] simulated-residual diagnostics for the observational model in Table 1 fitted as a beta regression on proportions transformed to the open interval (0, 1)[3] (top) and as an ordered beta regression on the untransformed proportions (bottom; used in the main analysis). Exact zeros and ones made up 56.5% and 3.6% of observations. Each panel shows a QQ plot of scaled residuals (left) and scaled residuals against model predictions (right). The departures under beta regression reflect the many plots with no $C_4$ cover, which the transformed response represents poorly.

**Supplementary Table 3.** Sensitivity of the best-supported observational model (Table 1) to adding mean annual precipitation (centre) and an exponential spatial covariance structure among site locations (right). Variance inflation factors in the Table 1 model were all below 2 (maximum 1.00). Adding mean annual precipitation changed AIC by -2.05. Moran's I on site-level residuals of the Table 1 model: I = -0.00, p = 0.849. Adding spatial covariance changed AIC by -4.8, with correlation decaying over an e-folding distance of about 246 km. Estimates are odds ratios.

| | **Table 1 model** | | | | **+ mean annual precipitation** | | | | **+ spatial covariance** | | | |
|---|---|---|---|---|---|---|---|---|---|---|---|---|
| *Predictors* | *Estimates* | *CI* | *Statistic* | *p* | *Estimates* | *CI* | *Statistic* | *p* | *Estimates* | *CI* | *Statistic* | *p* |
| Intercept | 0.002 | 0.001 – 0.006 | -10.441 | <0.001 | 0.002 | 0.001 – 0.006 | -10.630 | <0.001 | 0.002 | 0.000 – 0.010 | -7.805 | <0.001 |
| Proportion of PAR reaching the ground | 0.999 | 0.890 – 1.121 | -0.023 | 0.982 | 0.992 | 0.884 – 1.113 | -0.138 | 0.890 | 0.996 | 0.887 – 1.118 | -0.072 | 0.943 |
| Months ≥ 23 °C and ≥ 20 mm | 113.185 | 41.122 – 311.534 | 9.154 | <0.001 | 123.270 | 44.393 – 342.299 | 9.239 | <0.001 | 56.407 | 15.639 – 203.458 | 6.161 | <0.001 |
| Mean annual precipitation | | | | | 0.347 | 0.123 – 0.979 | -2.001 | 0.045 | | | | |
| N | $6_{block}$ | | | | $6_{block}$ | | | | $6_{block}$ | | | |
| | $112_{site\_code}$ | | | | $112_{site\_code}$ | | | | $112_{site\_code}$ | | | |
| | | | | | | | | | $1_{grp}$ | | | |
| Observations | 3186 | | | | 3186 | | | | 3186 | | | |
| AIC | 19.659 | | | | 17.607 | | | | 14.883 | | | |

**Supplementary Table 4.** Sensitivity of the best-supported observational model (Table 1) to sampling year: the main model (left), with sampling year as a covariate (centre), and refitted to the 44 sites surveyed in 2007–2008 (right). Estimates are odds ratios.

| | **Main** | | | | **+ sampling year** | | | | **Sites sampled 2007–2008** | | | |
|---|---|---|---|---|---|---|---|---|---|---|---|---|
| *Predictors* | *Estimates* | *CI* | *Statistic* | *p* | *Estimates* | *CI* | *Statistic* | *p* | *Estimates* | *CI* | *Statistic* | *p* |
| Intercept | 0.002 | 0.001 – 0.006 | -10.441 | <0.001 | 0.002 | 0.001 – 0.006 | -10.420 | <0.001 | 0.002 | 0.000 – 0.011 | -6.375 | <0.001 |
| Proportion of PAR reaching the ground | 0.999 | 0.890 – 1.121 | -0.023 | 0.982 | 0.999 | 0.890 – 1.121 | -0.017 | 0.987 | 1.032 | 0.847 – 1.258 | 0.311 | 0.756 |
| Months ≥ 23 °C and ≥ 20 mm | 113.185 | 41.122 – 311.534 | 9.154 | <0.001 | 112.280 | 40.771 – 309.206 | 9.134 | <0.001 | 161.105 | 26.963 – 962.609 | 5.572 | <0.001 |
| Sampling year | | | | | 0.850 | 0.348 – 2.075 | -0.357 | 0.721 | | | | |
| **Random Effects** | | | | | | | | | | | | |
| $\sigma^2$ | 9.79 | | | | 9.79 | | | | 10.13 | | | |
| $\tau_{00}$ | $0.27_{block:site\_code}$ | | | | $0.27_{block:site\_code}$ | | | | $0.28_{block:site\_code}$ | | | |
| | $20.64_{site\_code}$ | | | | $20.55_{site\_code}$ | | | | $21.22_{site\_code}$ | | | |
| ICC | 0.68 | | | | 0.68 | | | | 0.68 | | | |
| N | $6_{block}$ | | | | $6_{block}$ | | | | $6_{block}$ | | | |
| | $112_{site\_code}$ | | | | $112_{site\_code}$ | | | | $44_{site\_code}$ | | | |
| Observations | 3186 | | | | 3186 | | | | 1243 | | | |
| Marginal $R^2$ / Conditional $R^2$ | 0.422 / 0.816 | | | | 0.421 / 0.815 | | | | 0.415 / 0.813 | | | |

**Supplementary Table 5.** Proportional cover of $C_3$ grasses (left) and $C_3$ forbs (right), relative to total $C_3$ + $C_4$ cover, in relation to climate and ground-level PAR in the observational dataset (3186 plots), using the same predictors, random effects and model family (ordered beta) as the best-supported $C_4$ model (Table 1). Estimates are odds ratios per standard deviation.

| | **$C_3$ grass proportion** | | | | **$C_3$ forb proportion** | | | |
|---|---|---|---|---|---|---|---|---|
| *Predictors* | *Estimates* | *CI* | *Statistic* | *p* | *Estimates* | *CI* | *Statistic* | *p* |
| Intercept | 0.180 | 0.114 – 0.282 | -7.454 | <0.001 | 0.313 | 0.236 – 0.415 | -8.058 | <0.001 |
| Proportion of PAR reaching the ground | 0.829 | 0.772 – 0.890 | -5.185 | <0.001 | 1.270 | 1.190 – 1.356 | 7.157 | <0.001 |
| Months ≥ 23 °C and ≥ 20 mm | 0.184 | 0.115 – 0.292 | -7.170 | <0.001 | 0.713 | 0.538 – 0.945 | -2.350 | 0.019 |
| **Random Effects** | | | | | | | | |
| $\sigma^2$ | 0.42 | | | | 0.26 | | | |
| $\tau_{00}$ | 0.24 block:site_code | | | | 0.14 block:site_code | | | |
| | 5.36 site_code | | | | 2.20 site_code | | | |
| ICC | 0.93 | | | | 0.90 | | | |
| N | 6 block | | | | 6 block | | | |
| | 112 site_code | | | | 112 site_code | | | |
| Observations | 3186 | | | | 3186 | | | |
| Marginal $R^2$ / Conditional $R^2$ | 0.319 / 0.952 | | | | 0.069 / 0.906 | | | |

**Supplementary Table 6.** The crossover threshold compared with other climate predictors of $C_4$ proportional cover (ordered beta mixed models with ground-level PAR and site and block random intercepts). Top: AIC for models with a single climate predictor (the threshold count, mean annual temperature, maximum temperature of the warmest month, or the warm/cool-season rainfall ratio), with the four conventional WorldClim variables together (mean annual temperature, maximum temperature, precipitation seasonality and the rainfall ratio), and with the threshold added to the conventional variables. Bottom: estimates (odds ratios per standard deviation) for the threshold, rainfall-ratio, conventional-climate and combined models.

| Model | Climate predictors | AIC | ΔAIC |
|---|---|---|---|
| Threshold count (Table 1) | 1 | 19.7 | 23.6 |
| Mean annual temperature | 1 | 62.5 | 66.5 |
| Maximum temperature | 1 | 22.8 | 26.8 |
| Rainfall ratio | 1 | 68.7 | 72.7 |
| Conventional climate (4 variables) | 4 | -1.8 | 2.2 |
| Threshold + conventional climate | 5 | -4.0 | 0.0 |

| | Threshold (Table 1) | | | | Rainfall ratio | | | | Conventional climate | | | | Threshold + conventional climate | | | |
|---|---|---|---|---|---|---|---|---|---|---|---|---|---|---|---|---|
| *Predictors* | *Estimates* | *CI* | *Statistic* | *p* | *Estimates* | *CI* | *Statistic* | *p* | *Estimates* | *CI* | *Statistic* | *p* | *Estimates* | *CI* | *Statistic* | *p* |
| Intercept | 0.002 | 0.001 – 0.006 | -10.441 | <0.001 | 0.001 | 0.000 – 0.006 | -7.752 | <0.001 | 0.003 | 0.001 – 0.009 | -10.783 | <0.001 | 0.003 | 0.001 – 0.009 | -11.153 | <0.001 |
| Proportion of PAR reaching the ground | 0.999 | 0.890 – 1.121 | -0.023 | 0.982 | 0.991 | 0.883 – 1.113 | -0.150 | 0.881 | 0.998 | 0.889 – 1.119 | -0.038 | 0.970 | 1.000 | 0.891 – 1.122 | -0.006 | 0.995 |
| Months ≥ 23 °C and ≥ 20 mm | 113.185 | 41.122 – 311.534 | 9.154 | <0.001 | | | | | | | | | 4.198 | 1.058 – 16.660 | 2.040 | 0.041 |
| Warm/cool-season rainfall ratio | | | | | 51.711 | 12.477 – 214.317 | 5.439 | <0.001 | 13.873 | 5.283 – 36.428 | 5.339 | <0.001 | 7.420 | 2.576 – 21.369 | 3.714 | <0.001 |
| Mean annual temperature | | | | | | | | | 2.829 | 0.752 – 10.647 | 1.538 | 0.124 | 1.416 | 0.325 – 6.174 | 0.463 | 0.643 |
| Maximum temperature of the warmest month | | | | | | | | | 29.041 | 6.438 – 131.005 | 4.383 | <0.001 | 15.710 | 3.290 – 75.009 | 3.453 | 0.001 |
| Precipitation seasonality | | | | | | | | | 0.593 | 0.251 – 1.404 | -1.188 | 0.235 | 0.835 | 0.347 – 2.008 | -0.403 | 0.687 |
| N | $6_{\text{block}}$ | | | | $6_{\text{block}}$ | | | | $6_{\text{block}}$ | | | | $6_{\text{block}}$ | | | |
| | $112_{\text{site\_code}}$ | | | | $112_{\text{site\_code}}$ | | | | $112_{\text{site\_code}}$ | | | | $112_{\text{site\_code}}$ | | | |
| Observations | 3186 | | | | 3186 | | | | 3186 | | | | 3186 | | | |
| AIC | 19.659 | | | | 68.710 | | | | -1.781 | | | | -3.989 | | | |

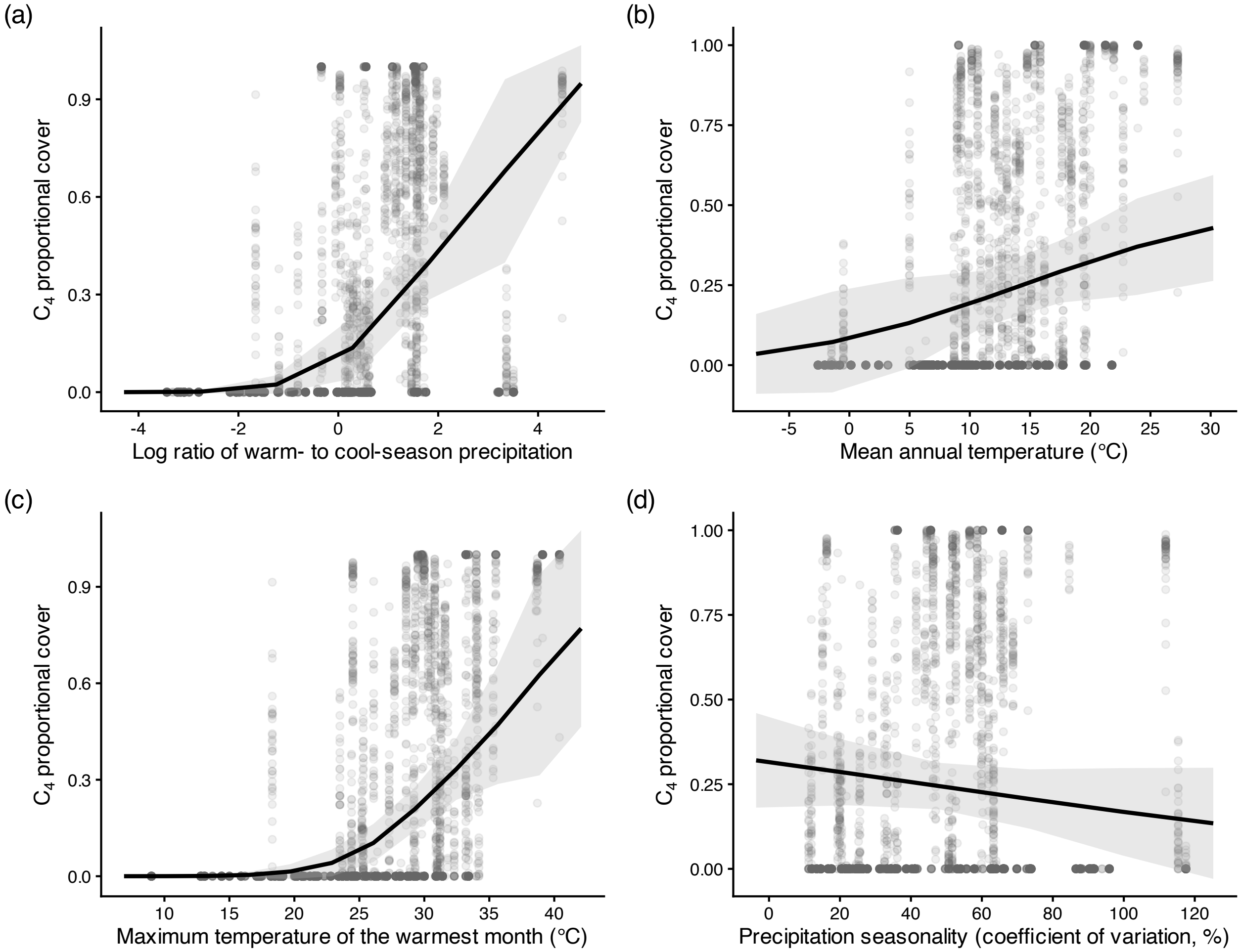


**Supplementary Fig. 5.** Predicted $C_4$ proportional cover in relation to the competing climate predictors in Supplementary Table 6: (a) the warm/cool-season rainfall ratio (rainfall-ratio model) and (b–d) mean annual temperature, maximum temperature of the warmest month and precipitation seasonality (conventional-climate model). Lines are predictions averaged over sites, with 95% confidence intervals; points are plots.

**Supplementary Table 7.** Number of sites contributing data at each plot age (years since treatments began) in the experimental models (Table 2, Figure 3; 48 sites in total).

| Age of plot (yrs) | Number of contributing sites |
|---:|---:|
| 1 | 48 |
| 2 | 47 |
| 3 | 41 |
| 4 | 34 |
| 5 | 30 |
| 6 | 27 |
| 7 | 23 |
| 8 | 20 |
| 9 | 16 |
| 10 | 14 |
| 11 | 14 |
| 12 | 12 |
| 13 | 7 |
| 14 | 8 |
| 15 | 4 |

**Supplementary Table 8.** Change in $C_4$ proportional cover per year of experiment for each treatment at low ($-1$ SD), mean and high (+1 SD) initial $C_4$ cover, from the ordered beta model with initial $C_4$ cover (Table 2), corresponding to the three panels of Figure 3. Values are odds ratios per year with 95% confidence intervals; values below 1 indicate a decline over time.

| Initial C4 cover | Treatment | Odds ratio per year [95% CI] | p |
|---|---|---|---|
| Low (-1 SD) | Control | 0.987 [0.950, 1.024] | 0.475 |
| Low (-1 SD) | Fence | 0.983 [0.943, 1.026] | 0.441 |
| Low (-1 SD) | NPK | 0.958 [0.919, 0.998] | 0.042 |
| Low (-1 SD) | NPK+Fence | 0.978 [0.932, 1.027] | 0.381 |
| Mean | Control | 0.974 [0.955, 0.994] | 0.010 |
| Mean | Fence | 0.958 [0.935, 0.982] | <0.001 |
| Mean | NPK | 0.924 [0.903, 0.945] | <0.001 |
| Mean | NPK+Fence | 0.897 [0.872, 0.923] | <0.001 |
| High (+1 SD) | Control | 0.962 [0.942, 0.981] | <0.001 |
| High (+1 SD) | Fence | 0.934 [0.906, 0.963] | <0.001 |
| High (+1 SD) | NPK | 0.891 [0.871, 0.912] | <0.001 |
| High (+1 SD) | NPK+Fence | 0.823 [0.794, 0.852] | <0.001 |

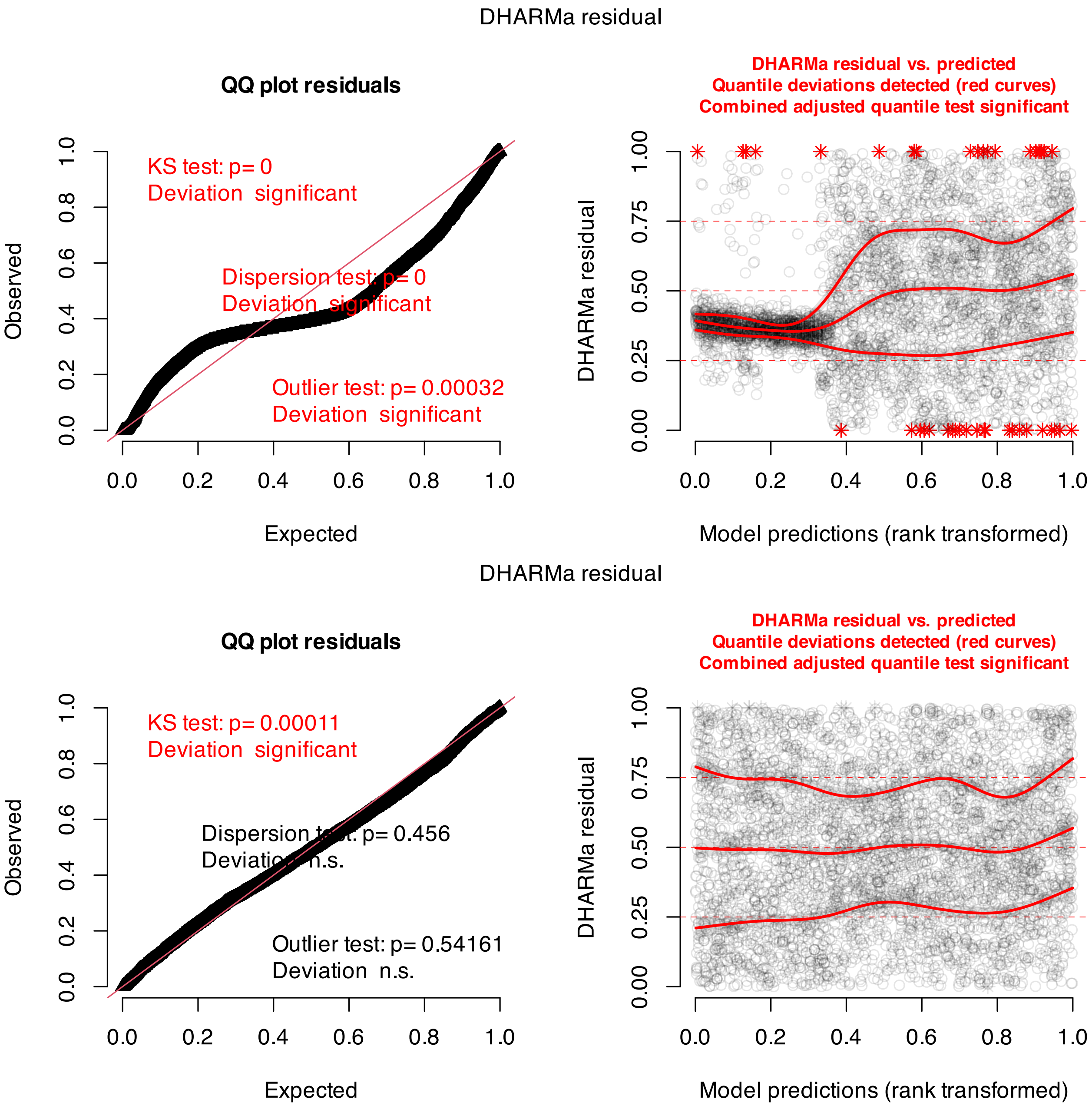


**Supplementary Fig. 6.** DHARMa simulated-residual diagnostics for the experimental model with initial $C_4$ cover (Table 2) fitted as a beta regression on proportions transformed to the open interval (0, 1) (top) and as an ordered beta regression on the untransformed proportions (bottom - used in the main analysis). Exact zeros and ones made up 39.5% and 1.8% of observations. Each panel shows a QQ plot of scaled residuals (left) and scaled residuals against model predictions (right).

**Supplementary Table 9.** Temporal trends in $C_4$ proportional cover in control plots in relation to calendar year (left) and calendar year interacting with the number of months above the climatic threshold (right), from ordered beta models with random intercepts for site and plot within site (no block as each block contains one control plot). Initial $C_4$ cover is expressed per standard deviation as in Table 2 in the main MS body. Calendar year and plot age were correlated across control plot-years (r = 0.59) because sites began treatments in different years.Trends with plot age are given in Supplementary Table 8. Estimates are odds ratios.

| | **Calendar year** | | | | **Calendar year × climate** | | | |
|---|---|---|---|---|---|---|---|---|
| *Predictors* | *Estimates* | *CI* | *Statistic* | *p* | *Estimates* | *CI* | *Statistic* | *p* |
| Intercept | 0.162 | 0.087 – 0.302 | -5.745 | <0.001 | 0.027 | 0.008 – 0.096 | -5.591 | <0.001 |
| Calendar year | 0.974 | 0.955 – 0.994 | -2.590 | 0.010 | 0.958 | 0.934 – 0.983 | -3.290 | 0.001 |
| Initial $C_4$ cover (per SD) | 16.714 | 8.808 – 31.717 | 8.616 | <0.001 | | | | |
| Calendar year × Initial $C_4$ cover (per SD) | 0.988 | 0.966 – 1.010 | -1.110 | 0.267 | | | | |
| Months ≥ 23 °C and ≥ 20 mm | | | | | 12.948 | 3.991 – 42.012 | 4.265 | <0.001 |
| Calendar year × Months ≥ 23 °C and ≥ 20 mm | | | | | 1.014 | 0.988 – 1.042 | 1.066 | 0.286 |
| **Random Effects** | | | | | | | | |
| $\sigma^2$ | 0.59 | | | | 0.60 | | | |
| $\tau_{00}$ | 0.37 $_{plot:site\_code}$ | | | | 0.35 $_{plot:site\_code}$ | | | |
| | 4.18 $_{site\_code}$ | | | | 9.95 $_{site\_code}$ | | | |
| ICC | 0.88 | | | | 0.95 | | | |
| N | 46 $_{plot}$ | | | | 46 $_{plot}$ | | | |
| | 48 $_{site\_code}$ | | | | 48 $_{site\_code}$ | | | |
| Observations | 1317 | | | | 1317 | | | |
| Marginal $R^2$ / Conditional $R^2$ | 0.604 / 0.954 | | | | 0.263 / 0.960 | | | |

**Supplementary Table 10.** Sensitivity of the treatment terms in the main experimental model (Table 2, main MS) to photosynthetic-pathway assignment (ordered beta models). Here we compared 1) as analysed (baseline); 2) with unassigned vascular taxa counted as $C_3$; 3) with genus-level $C_4$ assignments removed; 4) with all genus-level assignments ($C_3$ and $C_4$) removed; and 5) without the two sites with the largest share of unassigned cover (sandhill.us, 22.3%; sava.us, 22.1% of vascular cover). $C_4$ proportions and initial $C_4$ cover were recalculated for each version. Cells show odds ratios [95% CI] and asterisks denote where $p < 0.05$.

| Term | Baseline | Unassigned as C3 | No genus-level C4 | No genus-level assignments | Two sites dropped |
|---|---|---|---|---|---|
| Fence | 0.922 [0.733, 1.159] | 0.926 [0.733, 1.169] | 0.924 [0.732, 1.167] | 0.930 [0.728, 1.187] | 0.899 [0.713, 1.133] |
| NPK | 0.610 [0.499, 0.745]* | 0.613 [0.500, 0.752]* | 0.607 [0.495, 0.745]* | 0.607 [0.490, 0.752]* | 0.626 [0.511, 0.767]* |
| NPK+Fence | 0.534 [0.423, 0.675]* | 0.563 [0.444, 0.714]* | 0.531 [0.419, 0.674]* | 0.540 [0.421, 0.692]* | 0.535 [0.422, 0.677]* |
| Fence × plot age | 0.984 [0.954, 1.015] | 0.990 [0.959, 1.022] | 0.984 [0.954, 1.016] | 0.984 [0.954, 1.016] | 0.983 [0.953, 1.014] |
| NPK × plot age | 0.949 [0.921, 0.977]* | 0.942 [0.914, 0.971]* | 0.949 [0.921, 0.977]* | 0.939 [0.912, 0.968]* | 0.950 [0.922, 0.978]* |
| NPK+Fence × plot age | 0.921 [0.890, 0.954]* | 0.929 [0.897, 0.962]* | 0.919 [0.888, 0.952]* | 0.918 [0.887, 0.951]* | 0.921 [0.890, 0.954]* |
| Fence × initial C4 | 0.982 [0.771, 1.251] | 1.014 [0.793, 1.297] | 0.979 [0.766, 1.251] | 1.026 [0.804, 1.309] | 1.000 [0.783, 1.276] |
| NPK × initial C4 | 0.871 [0.699, 1.085] | 0.837 [0.670, 1.046] | 0.858 [0.687, 1.073] | 0.934 [0.753, 1.160] | 0.847 [0.679, 1.058] |
| NPK+Fence × initial C4 | 0.866 [0.678, 1.106] | 0.888 [0.692, 1.140] | 0.872 [0.680, 1.118] | 0.917 [0.717, 1.172] | 0.858 [0.670, 1.098] |
| Fence × plot age × initial C4 | 0.987 [0.953, 1.022] | 0.997 [0.963, 1.033] | 0.987 [0.953, 1.022] | 0.994 [0.962, 1.027] | 0.988 [0.954, 1.023] |
| NPK × plot age × initial C4 | 0.977 [0.945, 1.009] | 0.969 [0.937, 1.001] | 0.974 [0.942, 1.007] | 0.978 [0.949, 1.008] | 0.975 [0.944, 1.008] |
| NPK+Fence × plot age × initial C4 | 0.929 [0.894, 0.965]* | 0.938 [0.903, 0.975]* | 0.930 [0.895, 0.966]* | 0.941 [0.908, 0.975]* | 0.928 [0.893, 0.964]* |

**Supplementary Table 11.** Share of vascular plant cover excluded from the $C_4$ proportion (taxa with no assigned pathway, plus CAM and $C_3$–$C_4$ intermediate species) by treatment, across the crossover sites (years >= 1). Treatment differences were tested with an ordered beta mixed model with site, block and plot random intercepts (likelihood-ratio test: $x^2 = 28.67$, df = 9, $p < 0.001$). Across all sites and years, photosynthetic pathways were assigned at the species level for 85.1% of vascular plant cover and at the genus or family level for 13.8% (genus-level assignments made up 0.4% of $C_4$ cover). Note that 1.1% of vascular cover could not be assigned to a pathway and 0.1% belonged to CAM or $C_3$–$C_4$ intermediate species which were dropped.

| Treatment | Mean % of cover excluded | Plot-years |
|---|---|---|
| Control | 1.7 | 1317 |
| Fence | 1.0 | 826 |
| K | 1.3 | 1131 |
| N | 2.2 | 1129 |
| NK | 2.2 | 1133 |
| NP | 2.2 | 1131 |
| NPK | 2.0 | 1213 |
| NPK+Fence | 0.5 | 824 |
| P | 1.1 | 1132 |
| PK | 1.0 | 1132 |

**Supplementary Table 12.** Absolute (summed) cover of $C_4$ species (left) and $C_3$ species (right) in the experimental plots modelled with a Tweedie distribution (log link) and the same fixed and random effects as the main model shown in Table 2. Note that here, estimates are cover ratios.

| | $C_4$ cover | | | | $C_3$ cover | | | |
|---|---|---|---|---|---|---|---|---|
| *Predictors* | *Estimates* | *CI* | *Statistic* | *p* | *Estimates* | *CI* | *Statistic* | *p* |
| Intercept | 3.761 | 1.522 – 9.293 | 2.870 | 0.004 | 45.746 | 37.537 – 55.749 | 37.889 | <0.001 |
| Fence | 0.951 | 0.781 – 1.156 | -0.507 | 0.612 | 1.061 | 0.987 – 1.141 | 1.616 | 0.106 |
| NPK | 0.744 | 0.626 – 0.883 | -3.387 | 0.001 | 1.242 | 1.170 – 1.318 | 7.141 | <0.001 |
| NPK+Fence | 0.697 | 0.571 – 0.851 | -3.547 | <0.001 | 1.325 | 1.234 – 1.422 | 7.779 | <0.001 |
| Plot age | 0.980 | 0.961 – 0.998 | -2.157 | 0.031 | 1.002 | 0.995 – 1.008 | 0.487 | 0.626 |
| Initial $C_4$ cover (per SD) | 15.338 | 6.260 – 37.579 | 5.972 | <0.001 | 0.473 | 0.390 – 0.575 | -7.536 | <0.001 |
| Fence × Plot age | 0.979 | 0.950 – 1.008 | -1.425 | 0.154 | 1.002 | 0.991 – 1.013 | 0.343 | 0.732 |
| NPK × Plot age | 0.962 | 0.936 – 0.989 | -2.715 | 0.007 | 1.009 | 1.000 – 1.018 | 1.966 | 0.049 |
| NPK+Fence × Plot age | 0.943 | 0.912 – 0.974 | -3.555 | <0.001 | 1.017 | 1.006 – 1.028 | 3.153 | 0.002 |
| Fence × Initial $C_4$ cover (per SD) | 1.063 | 0.873 – 1.295 | 0.609 | 0.543 | 1.062 | 0.987 – 1.142 | 1.611 | 0.107 |
| NPK × Initial $C_4$ cover (per SD) | 0.910 | 0.759 – 1.090 | -1.024 | 0.306 | 1.096 | 1.030 – 1.167 | 2.874 | 0.004 |
| NPK+Fence × Initial $C_4$ cover (per SD) | 0.951 | 0.779 – 1.160 | -0.496 | 0.620 | 1.168 | 1.087 – 1.255 | 4.240 | <0.001 |
| Plot age × Initial $C_4$ cover (per SD) | 1.010 | 0.989 – 1.030 | 0.925 | 0.355 | 1.016 | 1.010 – 1.023 | 4.861 | <0.001 |
| Fence × Plot age × Initial $C_4$ cover (per SD) | 1.006 | 0.975 – 1.038 | 0.390 | 0.696 | 1.002 | 0.991 – 1.013 | 0.317 | 0.751 |
| NPK × Plot age × Initial $C_4$ cover (per SD) | 0.978 | 0.949 – 1.008 | -1.452 | 0.147 | 1.003 | 0.994 – 1.012 | 0.616 | 0.538 |
| NPK+Fence × Plot age × Initial $C_4$ cover (per SD) | 0.964 | 0.932 – 0.997 | -2.122 | 0.034 | 1.019 | 1.009 – 1.030 | 3.601 | <0.001 |
| **Random Effects** | | | | | | | | |
| $\sigma^2$ | 0.00 | | | | 0.11 | | | |
| $\tau_{00}$ | 0.29 $_{plot:block:site\_code}$ | | | | 0.04 $_{plot:block:site\_code}$ | | | |
| | 0.05 $_{block:site\_code}$ | | | | 0.03 $_{block:site\_code}$ | | | |
| | 8.97 $_{site\_code}$ | | | | 0.45 $_{site\_code}$ | | | |
| N | 53 $_{plot}$ | | | | 53 $_{plot}$ | | | |
| | 6 $_{block}$ | | | | 6 $_{block}$ | | | |
| | 48 $_{site\_code}$ | | | | 48 $_{site\_code}$ | | | |
| Observations | 4179 | | | | 4179 | | | |

**Supplementary Table 13.** Treatment effects on absolute $C_4$ and $C_3$ cover from the models in Supplementary Table 12. Annual change in cover within each treatment (% per year, at the mean initial $C_4$ cover) and the ratio of cover relative to control (at the mean plot age and mean initial $C_4$ cover).

| Response | Treatment | Annual change, % [95% CI] | Cover ratio vs control [95% CI] | p |
|---|---|---|---|---|
| C4 cover | Control | -2.0 [-3.9, -0.2] | – | – |
| C4 cover | Fence | -4.1 [-6.4, -1.9] | 0.951 [0.751, 1.204] | 0.892 |
| C4 cover | NPK | -5.8 [-7.8, -3.7] | 0.744 [0.605, 0.914] | 0.002 |
| C4 cover | NPK+Fence | -7.7 [-10.2, -5.1] | 0.697 [0.548, 0.886] | 0.001 |
| C3 cover | Control | 0.2 [-0.5, 0.8] | – | – |
| C3 cover | Fence | 0.4 [-0.6, 1.3] | 1.061 [0.973, 1.158] | 0.254 |
| C3 cover | NPK | 1.1 [0.4, 1.7] | 1.242 [1.156, 1.334] | <0.001 |
| C3 cover | NPK+Fence | 1.9 [1.0, 2.7] | 1.325 [1.217, 1.443] | <0.001 |

**Supplementary Table 14.** Proportion of photosynthetically active radiation (PAR) reaching the ground by treatment at the mean plot age, from a beta mixed model with treatment x plot age and site, block and plot random intercepts. Odds ratios compare each treatment with control (Dunnett-adjusted p as default via emmeans).

| Treatment | Proportion of PAR at ground [95% CI] | Odds ratio vs control [95% CI] | p |
|---|---|---|---|
| Control | 0.339 [0.265, 0.423] | – | – |
| Fence | 0.280 [0.213, 0.358] | 0.757 [0.653, 0.877] | <0.001 |
| NPK | 0.231 [0.173, 0.300] | 0.584 [0.512, 0.666] | <0.001 |
| NPK+Fence | 0.180 [0.133, 0.239] | 0.427 [0.367, 0.497] | <0.001 |

**Supplementary Table 15.** Factorial model of the change in $C_4$ proportional cover after five years (year 0 to year 4) with N, P and K and all their interactions, excluding fenced plots, at the subset of sites with the full factorial design. The N x P term tests a non additive relationship.

| | Change in $C_4$ proportion (year 0–4) | | | |
|---|---|---|---|---|
| *Predictors* | *Estimates* | *CI* | *Statistic* | *p* |
| Intercept | -0.048 | -0.114 – 0.019 | -1.416 | 0.157 |
| N | -0.065 | -0.120 – -0.010 | -2.332 | 0.020 |
| P | 0.043 | -0.012 – 0.098 | 1.536 | 0.125 |
| K | 0.049 | -0.005 – 0.104 | 1.775 | 0.076 |
| N × P | -0.062 | -0.141 – 0.017 | -1.544 | 0.123 |
| N × K | -0.032 | -0.111 – 0.046 | -0.807 | 0.420 |
| P × K | -0.085 | -0.163 – -0.006 | -2.112 | 0.035 |
| N × P × K | 0.057 | -0.055 – 0.168 | 0.999 | 0.318 |
| **Random Effects** | | | | |
| $\sigma^2$ | 0.04 | | | |
| $\tau_{00\ block:site\_code}$ | 0.00 | | | |
| $\tau_{00\ site\_code}$ | 0.02 | | | |
| ICC | 0.40 | | | |
| $N_{block}$ | 6 | | | |
| $N_{site\_code}$ | 27 | | | |
| Observations | 711 | | | |
| Marginal $R^2$ / Conditional $R^2$ | 0.045 / 0.425 | | | |

**Supplementary Table 16.** Pairwise comparisons between treatments in the change in $C_4$ proportional cover after five years (year 0 to year 4), from the model in Table 3. Estimates are differences (first treatment minus second). p-values are Tukey-adjusted.

| Contrast | Estimate | SE | df | t | p (Tukey) |
|---|---|---|---|---|---|
| Control – P | -0.043 | 0.029 | 746.1 | -1.48 | 0.901 |
| Control – K | -0.049 | 0.029 | 745.8 | -1.71 | 0.788 |
| Control – PK | -0.008 | 0.029 | 745.8 | -0.27 | 1.000 |
| Control – N | 0.066 | 0.029 | 746.5 | 2.27 | 0.408 |
| Control – NK | 0.048 | 0.029 | 745.8 | 1.68 | 0.804 |
| Control – NP | 0.085 | 0.029 | 745.8 | 2.94 | 0.096 |
| Control – NPK | 0.095 | 0.028 | 745.7 | 3.38 | 0.026 |
| Control – (NPK+Fence) | 0.094 | 0.032 | 751.4 | 2.93 | 0.100 |
| Control – Fence | 0.016 | 0.032 | 751.5 | 0.49 | 1.000 |
| P – K | -0.007 | 0.030 | 744.5 | -0.22 | 1.000 |
| P – PK | 0.035 | 0.030 | 744.5 | 1.18 | 0.976 |
| P – N | 0.108 | 0.030 | 745.6 | 3.63 | 0.011 |
| P – NK | 0.091 | 0.030 | 744.5 | 3.07 | 0.068 |
| P – NP | 0.127 | 0.030 | 744.5 | 4.29 | <0.001 |
| P – NPK | 0.138 | 0.029 | 745.1 | 4.72 | <0.001 |
| P – (NPK+Fence) | 0.136 | 0.033 | 748.6 | 4.17 | 0.001 |
| P – Fence | 0.058 | 0.033 | 749.1 | 1.78 | 0.748 |
| K – PK | 0.042 | 0.030 | 744.1 | 1.41 | 0.925 |
| K – N | 0.115 | 0.030 | 744.9 | 3.86 | 0.005 |
| K – NK | 0.098 | 0.030 | 744.1 | 3.30 | 0.034 |
| K – NP | 0.134 | 0.030 | 744.1 | 4.53 | <0.001 |
| K – NPK | 0.145 | 0.029 | 745.2 | 4.96 | <0.001 |
| K – (NPK+Fence) | 0.143 | 0.033 | 747.9 | 4.39 | <0.001 |
| K – Fence | 0.065 | 0.033 | 748.0 | 1.99 | 0.607 |
| PK – N | 0.073 | 0.030 | 744.9 | 2.47 | 0.288 |
| PK – NK | 0.056 | 0.030 | 744.1 | 1.90 | 0.672 |
| PK – NP | 0.092 | 0.030 | 744.1 | 3.12 | 0.059 |
| PK – NPK | 0.103 | 0.029 | 745.2 | 3.53 | 0.016 |
| PK – (NPK+Fence) | 0.101 | 0.033 | 747.9 | 3.11 | 0.060 |
| PK – Fence | 0.023 | 0.033 | 748.0 | 0.72 | 0.999 |
| N – NK | -0.017 | 0.030 | 744.9 | -0.58 | 1.000 |
| N – NP | 0.019 | 0.030 | 744.9 | 0.63 | 1.000 |
| N – NPK | 0.030 | 0.029 | 745.4 | 1.01 | 0.992 |
| N – (NPK+Fence) | 0.028 | 0.033 | 749.0 | 0.85 | 0.998 |
| N – Fence | -0.050 | 0.033 | 748.1 | -1.52 | 0.883 |
| NK – NP | 0.036 | 0.030 | 744.1 | 1.22 | 0.969 |
| NK – NPK | 0.047 | 0.029 | 745.2 | 1.61 | 0.843 |
| NK – (NPK+Fence) | 0.045 | 0.033 | 747.9 | 1.39 | 0.932 |
| NK – Fence | -0.033 | 0.033 | 748.0 | -1.00 | 0.992 |
| NP – NPK | 0.011 | 0.029 | 745.2 | 0.37 | 1.000 |
| NP – (NPK+Fence) | 0.009 | 0.033 | 747.9 | 0.27 | 1.000 |
| NP – Fence | -0.069 | 0.033 | 748.0 | -2.10 | 0.525 |
| NPK – (NPK+Fence) | -0.002 | 0.032 | 750.3 | -0.06 | 1.000 |
| NPK – Fence | -0.080 | 0.032 | 749.9 | -2.45 | 0.295 |
| (NPK+Fence) – Fence | -0.078 | 0.035 | 744.7 | -2.23 | 0.439 |

**Supplementary Table 17.** Change in $C_4$ proportional cover after five years in relation to treatment and pre-treatment soil phosphorus (log ppm P standardised; left 24 sites) and soil nitrogen (% N standardised; right 24 sites). Only one of the 27 sites lies in the tropics (latitude < 23.44°).

| | **Soil P** | | | | **Soil N** | | | |
|---|---|---|---|---|---|---|---|---|
| *Predictors* | *Estimates* | *CI* | *Statistic* | *p* | *Estimates* | *CI* | *Statistic* | *p* |
| Intercept | -0.039 | -0.106 – 0.027 | -1.156 | 0.248 | -0.038 | -0.105 – 0.030 | -1.092 | 0.275 |
| P | 0.045 | -0.015 – 0.104 | 1.471 | 0.142 | 0.047 | -0.013 – 0.107 | 1.554 | 0.121 |
| K | 0.053 | -0.006 – 0.113 | 1.765 | 0.078 | 0.056 | -0.004 – 0.115 | 1.825 | 0.068 |
| PK | 0.013 | -0.046 – 0.073 | 0.440 | 0.660 | 0.017 | -0.043 – 0.076 | 0.549 | 0.583 |
| N | -0.067 | -0.127 – -0.008 | -2.210 | 0.027 | -0.065 | -0.125 – -0.005 | -2.127 | 0.034 |
| NK | -0.057 | -0.116 – 0.002 | -1.884 | 0.060 | -0.056 | -0.116 – 0.004 | -1.845 | 0.065 |
| NP | -0.085 | -0.144 – -0.026 | -2.816 | 0.005 | -0.084 | -0.143 – -0.024 | -2.749 | 0.006 |
| NPK | -0.093 | -0.152 – -0.035 | -3.141 | 0.002 | -0.091 | -0.150 – -0.032 | -3.026 | 0.003 |
| NPK+Fence | -0.085 | -0.150 – -0.020 | -2.559 | 0.011 | -0.076 | -0.142 – -0.009 | -2.222 | 0.027 |
| Fence | -0.013 | -0.079 – 0.053 | -0.387 | 0.699 | -0.025 | -0.092 – 0.042 | -0.738 | 0.461 |
| Soil P | -0.020 | -0.086 – 0.045 | -0.609 | 0.543 | | | | |
| P × Soil P | 0.041 | -0.019 – 0.100 | 1.341 | 0.180 | | | | |
| K × Soil P | -0.008 | -0.067 – 0.051 | -0.266 | 0.790 | | | | |
| PK × Soil P | 0.034 | -0.025 – 0.093 | 1.128 | 0.260 | | | | |
| N × Soil P | -0.029 | -0.089 – 0.030 | -0.968 | 0.334 | | | | |
| NK × Soil P | -0.058 | -0.117 – 0.001 | -1.923 | 0.055 | | | | |
| NP × Soil P | -0.035 | -0.095 – 0.024 | -1.179 | 0.239 | | | | |
| NPK × Soil P | -0.021 | -0.079 – 0.037 | -0.708 | 0.479 | | | | |
| NPK+Fence × Soil P | -0.072 | -0.137 – -0.008 | -2.199 | 0.028 | | | | |
| Fence × Soil P | 0.019 | -0.047 – 0.084 | 0.565 | 0.572 | | | | |
| Soil N | | | | | 0.001 | -0.063 – 0.065 | 0.030 | 0.976 |
| P × Soil N | | | | | -0.061 | -0.118 – -0.003 | -2.060 | 0.040 |
| K × Soil N | | | | | -0.040 | -0.098 – 0.017 | -1.375 | 0.170 |
| PK × Soil N | | | | | -0.041 | -0.098 – 0.017 | -1.379 | 0.168 |
| N × Soil N | | | | | -0.015 | -0.072 – 0.043 | -0.495 | 0.621 |
| NK × Soil N | | | | | 0.008 | -0.049 – 0.066 | 0.285 | 0.776 |
| NP × Soil N | | | | | -0.039 | -0.096 – 0.019 | -1.318 | 0.188 |
| NPK × Soil N | | | | | -0.018 | -0.074 – 0.037 | -0.653 | 0.514 |
| NPK+Fence × Soil N | | | | | 0.021 | -0.056 – 0.099 | 0.540 | 0.589 |
| Fence × Soil N | | | | | -0.104 | -0.182 – -0.027 | -2.643 | 0.008 |
| **Random Effects** | | | | | | | | |
| $\sigma^2$ | 0.04 | | | | 0.04 | | | |
| $\tau_{00}$ | 0.00 $_{block:site\_code}$ | | | | 0.00 $_{block:site\_code}$ | | | |
| | 0.02 $_{site\_code}$ | | | | 0.02 $_{site\_code}$ | | | |
| ICC | 0.34 | | | | 0.34 | | | |
| N | 6 $_{block}$ | | | | 6 $_{block}$ | | | |
| | 24 $_{site\_code}$ | | | | 24 $_{site\_code}$ | | | |
| Observations | 766 | | | | 766 | | | |
| Marginal $R^2$ / Conditional $R^2$ | 0.081 / 0.394 | | | | 0.069 / 0.390 | | | |

To better understand whether the decline in $C_4$ observed above is accompanied by increases in $C_3$ grasses, $C_3$ forbs/herbs, or $C_3$ woody/shrub species, we fit three separate models (one per $C_3$ group) using the same `glmmTMB` temporal structure as the main experimental effects model. Each model is restricted to sites where that $C_3$ group was recorded at least once across the full experiment, with the logic being that sites with no forbs or no woody plants in their species pool cannot meaningfully contribute to a model of forb or woody cover change. $C_4$ grass is modelled across all crossover sites for direct comparison with the main model.

Zeroes are filled within sites (a plot where $C_3$ forbs were present at some point but absent in a given survey year receives a zero for that year) so that the full temporal trajectory is captured without dropping plots. The three $C_3$ models are presented alongside the main $C_4$ result to allow visual comparison, but because each draws on a different site subset, apparent offsetting patterns should be interpreted very cautiously. *That is, a $C_4$ decline and a $C_3$ grass increase may not be occurring at the same sites.*

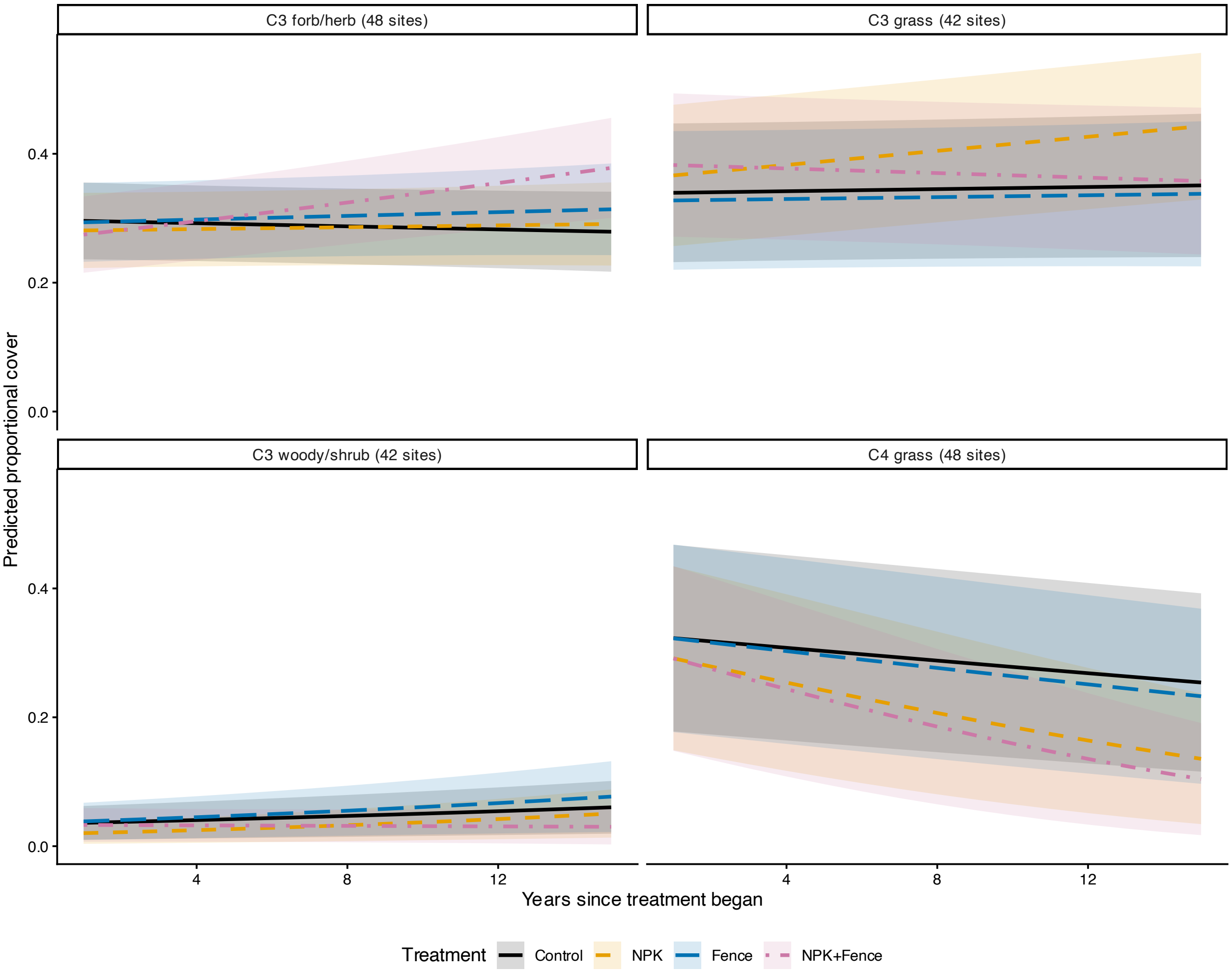


**Supplementary Fig. 7.** Temporal trajectories of proportional cover by functional group and treatment across the crossover-filtered experimental sites. $C_3$ grass, $C_3$ forb/herb, and $C_3$ woody/shrub are each restricted to sites where that group was recorded at least once across the experiment, so sample sizes differ between panels (shown in panels). Ribbons show 95% confidence intervals. As above, exercise caution interpreting compensatory effects here given the different subsamples of sites in each comparison.

**Supplementary Table 18.** Fixed-effect estimates (logit scale) from ordered beta `glmmTMB` temporal models per functional group. $C_4$ grass sample size matches the main model. $C_3$ groups are restricted to sites where that functional group was measured at some point during the experiment.

| Group | Sites | Plot-years | Term | Estimate | SE | CI low | CI high | p | p (BH) |
|---|---|---|---|---|---|---|---|---|---|
| C3 forb/herb | 48 | 4179 | Intercept | -1.221 | 0.179 | -1.571 | -0.871 | <0.001 | |
| C3 forb/herb | 48 | 4179 | Fence | -0.029 | 0.101 | -0.226 | 0.168 | 0.776 | |
| C3 forb/herb | 48 | 4179 | Fence × year | 0.015 | 0.012 | -0.007 | 0.038 | 0.180 | 0.297 |
| C3 forb/herb | 48 | 4179 | NPK | -0.099 | 0.090 | -0.276 | 0.077 | 0.269 | |
| C3 forb/herb | 48 | 4179 | NPK+Fence | -0.176 | 0.102 | -0.375 | 0.024 | 0.084 | |
| C3 forb/herb | 48 | 4179 | NPK+Fence × year | 0.048 | 0.012 | 0.026 | 0.071 | <0.001 | <0.001 |
| C3 forb/herb | 48 | 4179 | NPK × year | 0.011 | 0.010 | -0.009 | 0.031 | 0.267 | 0.356 |
| C3 forb/herb | 48 | 4179 | year | -0.007 | 0.007 | -0.021 | 0.007 | 0.311 | |
| C3 grass | 42 | 3839 | Intercept | -1.676 | 0.364 | -2.390 | -0.962 | <0.001 | |
| C3 grass | 42 | 3839 | Fence | -0.078 | 0.114 | -0.302 | 0.145 | 0.492 | |
| C3 grass | 42 | 3839 | Fence × year | -0.001 | 0.013 | -0.027 | 0.026 | 0.969 | 0.969 |
| C3 grass | 42 | 3839 | NPK | 0.147 | 0.104 | -0.056 | 0.350 | 0.155 | |
| C3 grass | 42 | 3839 | NPK+Fence | 0.298 | 0.114 | 0.075 | 0.521 | 0.009 | |
| C3 grass | 42 | 3839 | NPK+Fence × year | -0.017 | 0.013 | -0.043 | 0.009 | 0.198 | 0.297 |
| C3 grass | 42 | 3839 | NPK × year | 0.030 | 0.012 | 0.006 | 0.054 | 0.014 | 0.043 |
| C3 grass | 42 | 3839 | year | 0.005 | 0.008 | -0.011 | 0.022 | 0.525 | |
| C3 woody/shrub | 42 | 3730 | Intercept | -6.014 | 0.542 | -7.076 | -4.952 | <0.001 | |
| C3 woody/shrub | 42 | 3730 | Fence | 0.068 | 0.363 | -0.643 | 0.779 | 0.851 | |
| C3 woody/shrub | 42 | 3730 | Fence × year | 0.022 | 0.029 | -0.034 | 0.078 | 0.434 | 0.474 |
| C3 woody/shrub | 42 | 3730 | NPK | -0.856 | 0.320 | -1.484 | -0.228 | 0.008 | |
| C3 woody/shrub | 42 | 3730 | NPK+Fence | -0.071 | 0.377 | -0.810 | 0.667 | 0.850 | |
| C3 woody/shrub | 42 | 3730 | NPK+Fence × year | -0.065 | 0.031 | -0.127 | -0.003 | 0.039 | 0.093 |
| C3 woody/shrub | 42 | 3730 | NPK × year | 0.039 | 0.023 | -0.006 | 0.084 | 0.089 | 0.178 |
| C3 woody/shrub | 42 | 3730 | year | 0.056 | 0.014 | 0.029 | 0.083 | <0.001 | |
| C4 grass | 48 | 4179 | Intercept | -2.311 | 0.633 | -3.552 | -1.070 | <0.001 | |
| C4 grass | 48 | 4179 | Fence | 0.011 | 0.134 | -0.252 | 0.274 | 0.936 | |
| C4 grass | 48 | 4179 | Fence × year | -0.014 | 0.015 | -0.043 | 0.015 | 0.350 | 0.420 |
| C4 grass | 48 | 4179 | NPK | -0.202 | 0.117 | -0.431 | 0.027 | 0.084 | |
| C4 grass | 48 | 4179 | NPK+Fence | -0.174 | 0.138 | -0.444 | 0.096 | 0.206 | |
| C4 grass | 48 | 4179 | NPK+Fence × year | -0.098 | 0.017 | -0.131 | -0.065 | <0.001 | <0.001 |
| C4 grass | 48 | 4179 | NPK × year | -0.068 | 0.013 | -0.094 | -0.043 | <0.001 | <0.001 |
| C4 grass | 48 | 4179 | year | -0.043 | 0.009 | -0.060 | -0.026 | <0.001 | |

We partitioned the $C_4$ grass signal into independent $C_4$-origin lineages and fit a separate change model (year 0 to year 4) to each, using the same structure as the main change model (Table 3). Each lineage's model is restricted to sites where that lineage was recorded at least once across the experiment (a crude species-pool filter as with the functional group analyses above), with year-level zeroes filled within those sites where the lineage is not observed.

We only model four $C_4$ grass lineages due to sample size limitations in any others.

- **Andropogoneae** (tribe within Panicoideae NADP-ME)
- **Chloridoideae** (subfamily predominantly NAD-ME, with PCK)
- **MPC clade** (the Melinidinae, Panicinae and Cenchrinae radiation within the Paniceae, spanning all three biochemical subtypes of $C_4$)
- ***Aristida*** (a single genus forming its own $C_4$ lineage in the subfamily Aristidoideae[4])

Before fitting a change model to each lineage we assessed whether each lineage is actually modellable because the change model requires a plot to be present at both year 0 and year 4 (the paired requirement), and a plot that is zero at both timepoints contributes a change of exactly zero and is dropped. We therefore report for each lineage the number of plots and sites that survive a paired-and-non-zero requirement, together with how abundant the lineage is where it occurs and how zero-inflated it is across paired plots.

**Supplementary Table 19.** "Modellability" diagnostics for the four focal $C_4$ grass lineages across the crossover site pool. Paired plots were surveyed in both year 0 and year 4. "Modellable" plots and sites are the paired plots in which the lineage was present in at least one of those years. The percentage of paired plots with zero cover gives the share of paired plots in which the lineage was absent in both years. As in the main change model (Table 3), sites where $C_4$ plants were absent from all plots in both years are excluded. Median and maximum cover describe per plot abundance in modellable plots (mean of the two years). "Zero cover" - the % of plots with zero cover. "cov" - cover.

| C4 lineage | Modellable sites | M. plots | Paired plots | Zero cov. | Median cov. (%) | Max cov. (%) | Modelled |
|---|---|---|---|---|---|---|---|
| Andropogoneae | 18 | 391 | 816 | 52.1 | 31.1 | 152.5 | yes |
| Chloridoideae | 22 | 414 | 816 | 49.3 | 15.0 | 82.5 | yes |
| MPC clade | 13 | 188 | 816 | 77.0 | 7.5 | 65.0 | yes |
| Aristida | 8 | 87 | 816 | 89.3 | 2.0 | 52.5 | yes |

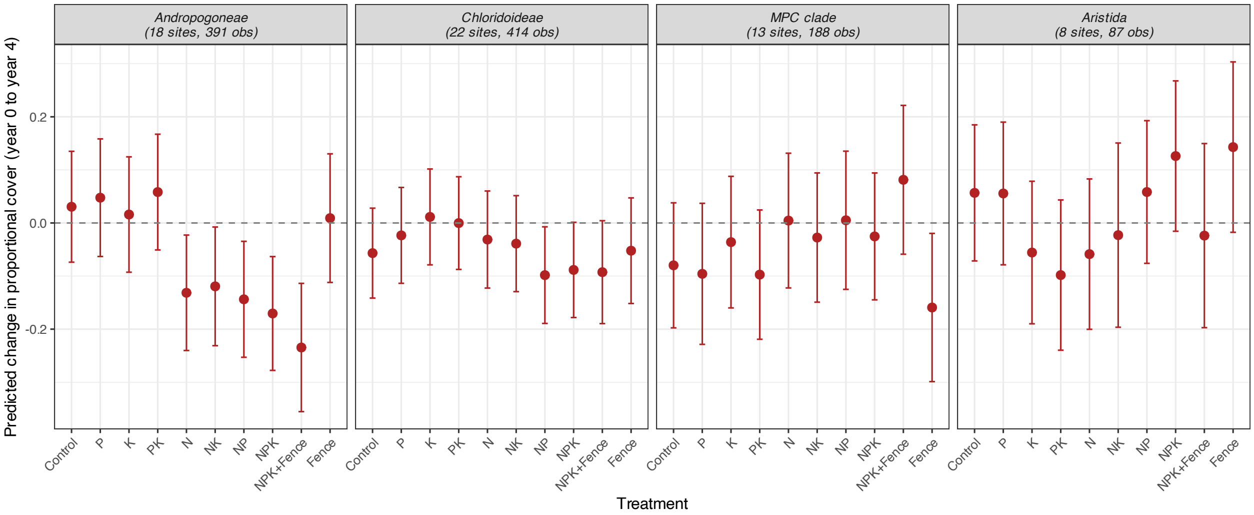


**Supplementary Fig. 8.** Predicted change in proportional cover (year 0 to year 4) for each focal $C_4$ grass lineage by treatment, from separate linear mixed models with the same structure as the main change model (Table 3). The four independent $C_4$ lineages (Andropogoneae, Chloridoideae, MPC clade and *Aristida*) are shown. Each uses only sites where it was recorded at least once with zeroes filled at those sites. Panel titles show the number of sites and plot-level observations.

**Supplementary Table 20.** Fixed-effect estimates from `lmer` change models for the four focal $C_4$ grass lineages, with sample sizes shown so the reliability of each estimate can be kept in mind. Each lineage's cover is expressed as a proportion of total live $C_3$ + $C_4$ cover and, as in the main change model (Table 3), proportions were rescaled into the open interval (0, 1) before the change between year 0 and year 4 was calculated[3]; the response is that change, modelled with a linear mixed model. Plots in which the lineage was absent in both years were excluded. Each lineage uses species-pool sites only, so cross-lineage effect-size comparisons should account for the differing site subsets.

| Lineage | Sites | Obs | Term | Estimate | SE | CI low | CI high | p | p (BH) |
|---|---|---|---|---|---|---|---|---|---|
| Andropogoneae | 18 | 391 | Intercept | 0.031 | 0.053 | -0.078 | 0.139 | 0.569 | |
| Andropogoneae | 18 | 391 | Fence | -0.021 | 0.052 | -0.124 | 0.081 | 0.681 | 0.834 |
| Andropogoneae | 18 | 391 | K | -0.015 | 0.044 | -0.102 | 0.073 | 0.741 | 0.834 |
| Andropogoneae | 18 | 391 | N | -0.162 | 0.044 | -0.249 | -0.075 | <0.001 | 0.003 |
| Andropogoneae | 18 | 391 | NK | -0.150 | 0.046 | -0.240 | -0.060 | 0.001 | 0.008 |
| Andropogoneae | 18 | 391 | NP | -0.174 | 0.045 | -0.262 | -0.087 | <0.001 | 0.001 |
| Andropogoneae | 18 | 391 | NPK | -0.201 | 0.043 | -0.286 | -0.116 | <0.001 | <0.001 |
| Andropogoneae | 18 | 391 | NPK+Fence | -0.265 | 0.052 | -0.367 | -0.163 | <0.001 | <0.001 |
| Andropogoneae | 18 | 391 | P | 0.017 | 0.045 | -0.072 | 0.106 | 0.708 | 0.834 |
| Andropogoneae | 18 | 391 | PK | 0.028 | 0.044 | -0.060 | 0.115 | 0.535 | 0.713 |
| Chloridoideae | 22 | 414 | Intercept | -0.057 | 0.043 | -0.145 | 0.031 | 0.198 | |
| Chloridoideae | 22 | 414 | Fence | 0.005 | 0.045 | -0.084 | 0.093 | 0.919 | 0.973 |
| Chloridoideae | 22 | 414 | K | 0.068 | 0.039 | -0.008 | 0.145 | 0.080 | 0.412 |
| Chloridoideae | 22 | 414 | N | 0.026 | 0.039 | -0.052 | 0.103 | 0.515 | 0.713 |
| Chloridoideae | 22 | 414 | NK | 0.018 | 0.039 | -0.059 | 0.094 | 0.646 | 0.831 |
| Chloridoideae | 22 | 414 | NP | -0.041 | 0.039 | -0.118 | 0.035 | 0.290 | 0.634 |
| Chloridoideae | 22 | 414 | NPK | -0.032 | 0.038 | -0.107 | 0.044 | 0.411 | 0.676 |
| Chloridoideae | 22 | 414 | NPK+Fence | -0.036 | 0.044 | -0.122 | 0.050 | 0.413 | 0.676 |
| Chloridoideae | 22 | 414 | P | 0.033 | 0.039 | -0.043 | 0.110 | 0.389 | 0.676 |
| Chloridoideae | 22 | 414 | PK | 0.057 | 0.038 | -0.017 | 0.130 | 0.132 | 0.432 |
| MPC clade | 13 | 188 | Intercept | -0.080 | 0.060 | -0.214 | 0.054 | 0.213 | |
| MPC clade | 13 | 188 | Fence | -0.079 | 0.061 | -0.200 | 0.041 | 0.196 | 0.587 |
| MPC clade | 13 | 188 | K | 0.044 | 0.051 | -0.056 | 0.144 | 0.390 | 0.676 |
| MPC clade | 13 | 188 | N | 0.084 | 0.052 | -0.019 | 0.188 | 0.109 | 0.432 |
| MPC clade | 13 | 188 | NK | 0.052 | 0.050 | -0.047 | 0.152 | 0.299 | 0.634 |
| MPC clade | 13 | 188 | NP | 0.085 | 0.054 | -0.023 | 0.192 | 0.121 | 0.432 |
| MPC clade | 13 | 188 | NPK | 0.054 | 0.048 | -0.040 | 0.149 | 0.256 | 0.615 |
| MPC clade | 13 | 188 | NPK+Fence | 0.161 | 0.061 | 0.041 | 0.281 | 0.009 | 0.054 |
| MPC clade | 13 | 188 | P | -0.016 | 0.055 | -0.125 | 0.093 | 0.773 | 0.843 |
| MPC clade | 13 | 188 | PK | -0.018 | 0.051 | -0.117 | 0.082 | 0.729 | 0.834 |
| Aristida | 8 | 87 | Intercept | 0.057 | 0.064 | -0.072 | 0.185 | 0.381 | |
| Aristida | 8 | 87 | Fence | 0.086 | 0.103 | -0.118 | 0.291 | 0.404 | 0.676 |
| Aristida | 8 | 87 | K | -0.112 | 0.093 | -0.297 | 0.072 | 0.229 | 0.591 |
| Aristida | 8 | 87 | N | -0.115 | 0.095 | -0.305 | 0.075 | 0.230 | 0.591 |
| Aristida | 8 | 87 | NK | -0.080 | 0.108 | -0.294 | 0.135 | 0.462 | 0.676 |
| Aristida | 8 | 87 | NP | 0.002 | 0.093 | -0.183 | 0.186 | 0.986 | 0.990 |
| Aristida | 8 | 87 | NPK | 0.069 | 0.095 | -0.121 | 0.259 | 0.470 | 0.676 |
| Aristida | 8 | 87 | NPK+Fence | -0.081 | 0.108 | -0.295 | 0.134 | 0.457 | 0.676 |
| Aristida | 8 | 87 | P | -0.001 | 0.093 | -0.186 | 0.184 | 0.990 | 0.990 |
| Aristida | 8 | 87 | PK | -0.155 | 0.095 | -0.345 | 0.035 | 0.109 | 0.432 |

**Supplementary Table 21.** Change in proportional cover (year 0 to year 4) of all $C_4$ species, all $C_4$ grasses and $C_4$ forbs by treatment, from linear mixed models with the same structure as the lineage models (Supplementary Table 20). Cells show estimates [95% CI] and p-values; column headers give the number of sites and plot-level observations.

| Term | All C4 (26 sites, 678 obs) | All C4 grass (26 sites, 668 obs) | C4 forb (8 sites, 94 obs) |
|---|---|---|---|
| Intercept | -0.036 [-0.117, 0.045], p = 0.376 | -0.037 [-0.113, 0.040], p = 0.342 | 0.091 [-0.082, 0.264], p = 0.248 |
| P | 0.051 [-0.016, 0.119], p = 0.135 | 0.026 [-0.044, 0.096], p = 0.460 | 0.007 [-0.009, 0.022], p = 0.396 |
| K | 0.056 [-0.010, 0.122], p = 0.096 | 0.033 [-0.036, 0.102], p = 0.346 | 0.012 [-0.002, 0.025], p = 0.085 |
| PK | 0.026 [-0.041, 0.093], p = 0.451 | 0.011 [-0.058, 0.080], p = 0.747 | 0.007 [-0.006, 0.020], p = 0.296 |
| N | -0.071 [-0.138, -0.004], p = 0.038 | -0.074 [-0.143, -0.005], p = 0.036 | 0.012 [-0.004, 0.027], p = 0.140 |
| NK | -0.042 [-0.109, 0.025], p = 0.220 | -0.049 [-0.118, 0.020], p = 0.163 | 0.010 [-0.005, 0.024], p = 0.178 |
| NP | -0.091 [-0.158, -0.024], p = 0.008 | -0.108 [-0.177, -0.040], p = 0.002 | 0.019 [0.005, 0.033], p = 0.009 |
| NPK | -0.104 [-0.169, -0.039], p = 0.002 | -0.117 [-0.184, -0.049], p < 0.001 | 0.019 [0.004, 0.034], p = 0.015 |
| NPK+Fence | -0.101 [-0.176, -0.025], p = 0.009 | -0.127 [-0.206, -0.048], p = 0.002 | 0.015 [-0.002, 0.033], p = 0.086 |
| Fence | -0.016 [-0.092, 0.060], p = 0.688 | -0.010 [-0.089, 0.068], p = 0.799 | 0.013 [-0.013, 0.039], p = 0.318 |

**Supplementary Table 22.** Likelihood-ratio test of a lineage x treatment interaction in a stacked change model (year 0 to year 4) including the four focal $C_4$ grass lineages, at the sites where at least two lineages co-occurred. Model includes site, block and plot random intercepts.

| Model | npar | AIC | logLik | Chisq | Df | Pr(>Chisq) |
|---|---|---|---|---|---|---|
| Treatment + lineage | 17 | 23.413 | 5.293 | | | |
| Treatment × lineage | 44 | 4.340 | 41.830 | 73.074 | 27 | <0.001 |

**Supplementary Table 23.** Sensitivity of the treatment terms in the main experimental model (Table 2, model 2) to site selection: as analysed (48 sites with at least two months above the crossover threshold), excluding sites with no $C_4$ cover in any analysed plot-year (39 sites), and excluding sites with mean pre-treatment $C_4$ proportional cover below 0.05 or above 0.95 (33 sites). Plot age and initial $C_4$ cover were re-centred within each subset, with initial cover scaled by the same standard deviation as in Table 2. Cells show odds ratios [95% CI]; asterisks denote $p < 0.05$.

| Term | As analysed | C4 present | Initial C4 0.05–0.95 |
|---|---|---|---|
| Fence | 0.922 [0.733, 1.159] | 0.921 [0.739, 1.149] | 1.015 [0.812, 1.269] |
| NPK | 0.610 [0.499, 0.745]* | 0.590 [0.489, 0.712]* | 0.534 [0.443, 0.644]* |
| NPK+Fence | 0.534 [0.423, 0.675]* | 0.519 [0.415, 0.650]* | 0.474 [0.377, 0.596]* |
| Fence × plot age | 0.984 [0.954, 1.015] | 0.980 [0.952, 1.009] | 0.981 [0.952, 1.010] |
| NPK × plot age | 0.949 [0.921, 0.977]* | 0.943 [0.918, 0.967]* | 0.929 [0.905, 0.954]* |
| NPK+Fence × plot age | 0.921 [0.890, 0.954]* | 0.903 [0.874, 0.932]* | 0.885 [0.856, 0.915]* |
| Fence × initial C4 | 0.982 [0.771, 1.251] | 1.002 [0.787, 1.277] | 0.955 [0.728, 1.253] |
| NPK × initial C4 | 0.871 [0.699, 1.085] | 0.886 [0.711, 1.105] | 0.997 [0.782, 1.271] |
| NPK+Fence × initial C4 | 0.866 [0.678, 1.106] | 0.895 [0.701, 1.143] | 1.103 [0.835, 1.457] |
| Fence × plot age × initial C4 | 0.987 [0.953, 1.022] | 0.987 [0.954, 1.022] | 0.981 [0.946, 1.019] |
| NPK × plot age × initial C4 | 0.977 [0.945, 1.009] | 0.977 [0.945, 1.010] | 1.001 [0.966, 1.038] |
| NPK+Fence × plot age × initial C4 | 0.929 [0.894, 0.965]* | 0.930 [0.895, 0.966]* | 0.950 [0.910, 0.991]* |

### Supplement References


1. Luo, X. et al. Mapping the global distribution of $C_4$ vegetation using observations and optimality theory. Nat. Commun. 15, 1219 (2024).
2. Hartig, F., Lohse, L., Leite, M. de S. & Werneke, C. DHARMa: residual diagnostics for hierarchical (multi-level/mixed) regression models. R package (2026).
3. Smithson, M. & Verkuilen, J. A better lemon squeezer? Maximum-likelihood regression with beta-distributed dependent variables. Psychol. Methods 11, 54–71 (2006).

4. Grass Phylogeny Working Group II. New grass phylogeny resolves deep evolutionary relationships and discovers $C_4$ origins. New Phytol. 193, 304–312 (2012).
